# Deep immunophenotyping reveals circulating activated lymphocytes in individuals at risk for rheumatoid arthritis

**DOI:** 10.1101/2023.07.03.547507

**Authors:** Jun Inamo, Joshua Keegan, Alec Griffith, Tusharkanti Ghosh, Alice Horisberger, Kaitlyn Howard, John Pulford, Ekaterina Murzin, Brandon Hancock, Anna Helena Jonsson, Jennifer Seifert, Marie L. Feser, Jill M. Norris, Ye Cao, William Apruzzese, S. Louis Bridges, Vivian Bykerk, Susan Goodman, Laura Donlin, Gary S. Firestein, Harris Perlman, Joan M. Bathon, Laura B. Hughes, Darren Tabechian, Andrew Filer, Costantino Pitzalis, Jennifer H. Anolik, Larry Moreland, Joel M. Guthridge, Judith A. James, Michael B. Brenner, Soumya Raychaudhuri, Jeffrey A. Sparks, The Accelerating Medicines Partnership RA/SLE Network, V. Michael Holers, Kevin D. Deane, James A. Lederer, Deepak A. Rao, Fan Zhang

## Abstract

Rheumatoid arthritis (RA) is a systemic autoimmune disease with currently no universally highly effective prevention strategies. Identifying pathogenic immune phenotypes in ‘At-Risk’ populations prior to clinical disease onset is crucial to establishing effective prevention strategies. Here, we applied mass cytometry to deeply characterize the immunophenotypes in blood from At-Risk individuals identified through the presence of serum antibodies to citrullinated protein antigens (ACPA) and/or first-degree relative (FDR) status (n=52), as compared to established RA (n=67), and healthy controls (n=48). We identified significant cell expansions in At-Risk individuals compared with controls, including CCR2+CD4+ T cells, T peripheral helper (Tph) cells, type 1 T helper cells, and CXCR5+CD8+ T cells. We also found that CD15+ classical monocytes were specifically expanded in ACPA-negative FDRs, and an activated PAX5^low^ naïve B cell population was expanded in ACPA-positive FDRs. Further, we developed an “RA immunophenotype score” classification method based on the degree of enrichment of cell states relevant to established RA patients. This score significantly distinguished At-Risk individuals from controls. In all, we systematically identified activated lymphocyte phenotypes in At-Risk individuals, along with immunophenotypic differences among both ACPA+ and ACPA-FDR At-Risk subpopulations. Our classification model provides a promising approach for understanding RA pathogenesis with the goal to further improve prevention strategies and identify novel therapeutic targets.

## Introduction

Rheumatoid arthritis (RA) is a prototypical autoimmune disease that affects 0.8-1.0% of the population^1, 2^. Although the presence of inflammatory arthritis is the hallmark of clinical RA, diagnosis and treatments are usually delayed, and no cure has been found^3^. Recent studies have shown that in seropositive RA there is a prolonged pre-RA phase that is characterized by blood elevations of biomarkers^4–6^, including antibodies to citrullinated protein antigens (ACPA), prior to the onset of ‘clinical RA’ (e.g., the first appearance of inflammatory arthritis)^7, 8^.

Notably, individuals can be defined for studies as being at high risk for future RA (e.g., At-Risk) due to being a first-degree relative (FDR) of a patient with RA and/or having elevations of ACPA in the peripheral blood^9–12^. In particular, serum elevations of ACPA are highly predictive of the future development of RA^13–16^, and multiple studies of ACPA+ At-Risk individuals have been performed to evaluate possible preventive strategies. However, trials of corticosteroids, atorvastatin, methotrexate, and B-cell depleted therapy demonstrated no significant effect of these agents in preventing progression to clinical RA^17–20^.

The mechanisms that drive autoimmunity in At-Risk states are still unclear, but likely involve a complex interplay between genetics, environmental factors, mucosal endotypes, and immunophenotypes^21–25^. Thus, investigating the spectrum of molecular and cellular changes in individuals who are in the At-Risk state is key to identifying predictive markers and phenotypes to further develop accurate prediction models for future RA, and identify targets for preventive interventions^1, 21, 26^. There are an increasing number of studies focused on characterizing immunophenotypes and biomarkers in At-Risk individuals^27–32^. We reasoned that applying high-dimensional cytometric immunophenotyping combined with computational integration strategies in a systematic manner would reveal additional relevant features in RA immune activation that would inform the understanding of disease pathogenesis.

Multi-institutional studies using single-cell technologies are advancing our understanding of autoimmune disease heterogeneity, in part through the increased power of uniform analyses of pooled cohorts^33–36^. Our recent work of harmonizing single-cell transcriptomics, mass cytometry, and single-cell multimodal CITE-seq from multiple clinical sites has uncovered key immune cell populations in inflamed synovium from patients with established RA^33, 37^. We and others have developed robust single-cell integration methods and identified specific RA-relevant immune populations^34, 38–42^, including our recently identified GZMK+ CD8 T cells^43^, previously identified T peripheral helper (Tph) and T follicular helper (Tfh) cells^33, 44^, pro-inflammatory myeloid cells like IL1B+ HBEGF+^33, 45^, IFN- and TNF-driven CXCL10+CCL2+ macrophages^34^, and NR4A+ B cells^46^. However, whether any of these pathogenic populations in the inflamed tissues are already present and altered in the circulation during the pre-RA phase of disease is unclear^34, 38–42^. Such populations may represent key treatment targets to prevent development of clinical RA.

Under the SERA (Studies of the Etiologies of Rheumatoid Arthritis) umbrella, as well as through the AMP RA/SLE (Accelerating Medicines Partnership Rheumatoid Arthritis/Systemic Lupus Erythematosus) Network, multi-institutional cohorts of At-Risk individuals, established RA, and relevant controls have been established to study the natural history of RA^27–32^. Here, we performed deep single-cell immunophenotyping using large mass cytometry data to characterize peripheral blood mononuclear cells (PBMCs) alterations in At-Risk individuals compared with established RA and healthy controls. Using our integrative and classification strategies, we identified multiple immune cell populations expanded in blood of various At-Risk populations based on the presence of ACPA and/or FDR status. Further, we built an “RA immunophenotype score” classification model and demonstrated its efficacy in quantifying and differentiating established RA, At-Risk individuals, and healthy controls. Our computational strategies and immunophenotypic findings define specific features of immune dysregulation in preclinical RA and nominate new potential targets for immunophenotype-based preventive strategies.

## Results

### Collection of peripheral blood samples from individuals with RA, At-Risk RA, and controls

We utilized PBMCs from individuals at risk for RA (At-Risk RA, n=52) and established RA (n=67) who were enrolled in the AMP RA/SLE Network. At-Risk individuals were sub-categorized based on their family history and/or the positivity of ACPA into FDR+ACPA- (n=23), FDR-ACPA+ (n=9), and FDR+ACPA+ (n=20) (**Figure 1A**, **Supplementary Table 1**). Similarly, RA patients were categorized into ACPA+ and ACPA-. For comparison, we collected PBMCs from healthy individuals as controls (n=48). All the collected PBMC samples (n=167) with other consortium samples (Systemic lupus erythematosus (SLE), n=140) were randomly distributed based on disease status, clinical site, and sex into 23 technical batches to minimize effects from site differences and other demographics. We then applied mass cytometry in one central site, so that all the antibody staining and preprocessing were done uniformly in one place (**Methods**).

Next, we applied computational integrative and association algorithms to identify unique co-varying phenotypical changes across different preclinical and clinical individual groups (**Figure 1A**). We applied an optimized downsampling strategy to analyze all mononuclear cells as well as specific immune cell lineages for computational efficiency (**Methods**). We next performed a sensitivity analysis by changing the downsampling proportions and confirmed that the immune cell clusters detected by different downsampling parameters are stable (**Extended Data Fig.1**). In total, we analyzed 1,640,747 cells for all mononuclear cells analysis (167 individuals), and 2,196,578 T cells (163 individuals), 1,886,084 myeloid cells (161 individuals), 1,918,711 B cells (167 individuals), and 2,008,997 NK cells (160 individuals) for each cell type analysis. To correct the technical batch effect and inter-individual variation, we applied a single-cell batch effect correction algorithm^38^ and quantified the improvement of mixture levels across technical batches, clinical sites, and individual samples after correction^34, 38^ (**Methods**). After batch effect correction, the degree of mixing levels across batches, race, and sites was significantly increased compared to before correction (**Extended Data Fig.2**). For accurate integration, we confirmed that the mixing levels for cell type, measured by LISI (Local Inverse Simpson’s Index)^38, 47^, as equal to 1, reflecting a correct separation of unique cell types throughout the integrative embedding.

### Single-cell proteomic profiling of mononuclear cells defines differential immune cell abundance in At-Risk RA

We first investigated all mononuclear cells from 167 individuals to characterize the landscape of immune cell heterogeneity, and then quantified the altered cell abundance across differential clinical phenotypes. We defined four major immune cell types, including T cells, myeloid cells, B cells, and NK cells, based on canonical protein markers and projected them into low-dimensional space (**Fig.1B**). To identify significant shifts of major cell type abundance between At-Risk RA and controls, we applied complementary computational strategies, including cluster-based mixed-effects modeling of associations of single cells (MASC)^40^ accounting for covariates age and sex, and cluster-free covarying neighborhood analysis (CNA)^41^ that identifies dominant co-vary cell type abundance accounting for covariates age and sex. We observed significantly expanded myeloid cells in At-Risk RA (*p* = 0.015, odds ratio (OR) = 1.28)) (**Fig. 1C**). Conversely, CD8 T cells (OR = 0.83) and NK cells (OR = 0.81) were depleted in At-Risk RA compared with controls. We also observed heterogeneity within CD4 T cells and their variable association levels, which suggested a need for fine-grained cell type-specific analysis.

**Fig.1:**
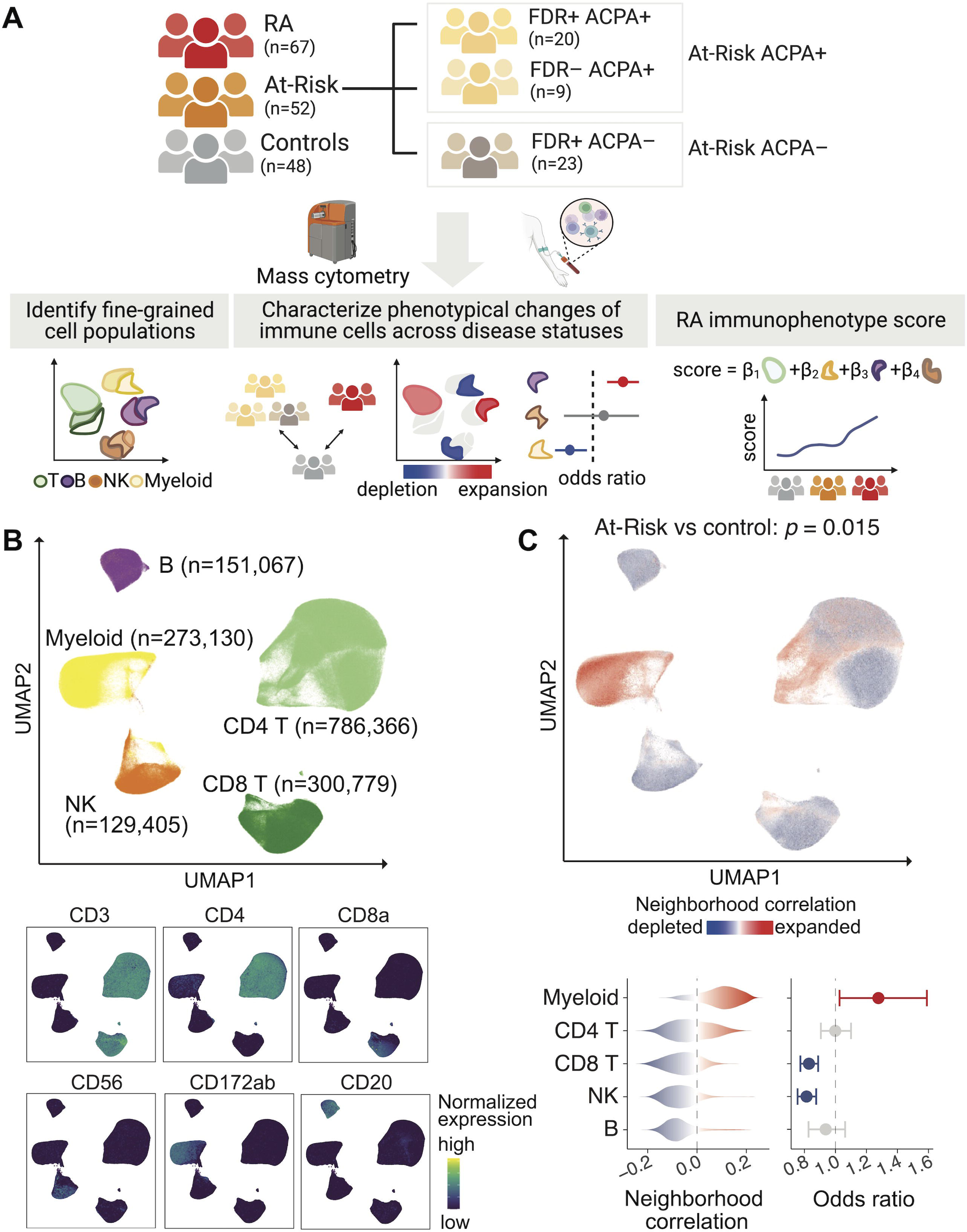
Overview of mass cytometry pipeline and cell type abundance analysis for At-Risk RA individuals using all mononuclear cells. **A.** Description of study design regarding patient recruitment, clinical classification, and computational strategies, **B.** Identified major immune cell types among all mononuclear cells and canonical protein expression in UMAP, **C.** At-Risk associations compared to controls for all mononuclear cells. P-value is generated from co-varying neighborhood (CNA) analysis. Cells in UMAP are colored for expansion (red) or depletion (blue) in At-Risk RA. For each cell type, distributions of At-Risk associated cell neighborhood correlations and odds ratios with 95% confidence intervals are shown. All the At-Risk association testing are adjusted for age and sex.

### Cell type-specific analysis reveals fine-grained cell subpopulations

For each major immune cell type, we defined fine-grained cell states based on expression of 48 protein markers and quantified cluster abundances and phenotypical changes (**Fig.2**, **Extended Data Fig.2**, **Supplementary Table 2**). We identified 79 immune cell clusters in total, including 26 T cell clusters, 16 myeloid clusters, 20 B cell clusters, and 17 NK cell clusters. We described differentially expressed proteins for each cluster and summarized the statistics (**Supplementary Table 3**). The T cells clustered broadly into CD4 and CD8 T cell subsets (**Fig.2A**, **Extended Data Fig.3**). Among the CD4+ T cells, naïve T cells (T-0, T-2) separated from memory cell populations, and a distinct population of regulatory T cells (T-9) marked by FoxP3 and CD25 clustered separately. The memory CD4 T cells included CXCR5+ Tfh cells (T-7, T-11) as well as a cluster of Tph cells (T-14), a B cell-helper population highly enriched in RA joints^33, 44^. CD8 T cells also separated in naïve and effector/memory subsets, with the memory cells segregating into distinct Granzyme B+ and Granzyme K+ subpopulations (T-3, T-8, T-15), as was also observed among CD8 T cells in RA synovium^43, 48^.

**Fig.2:**
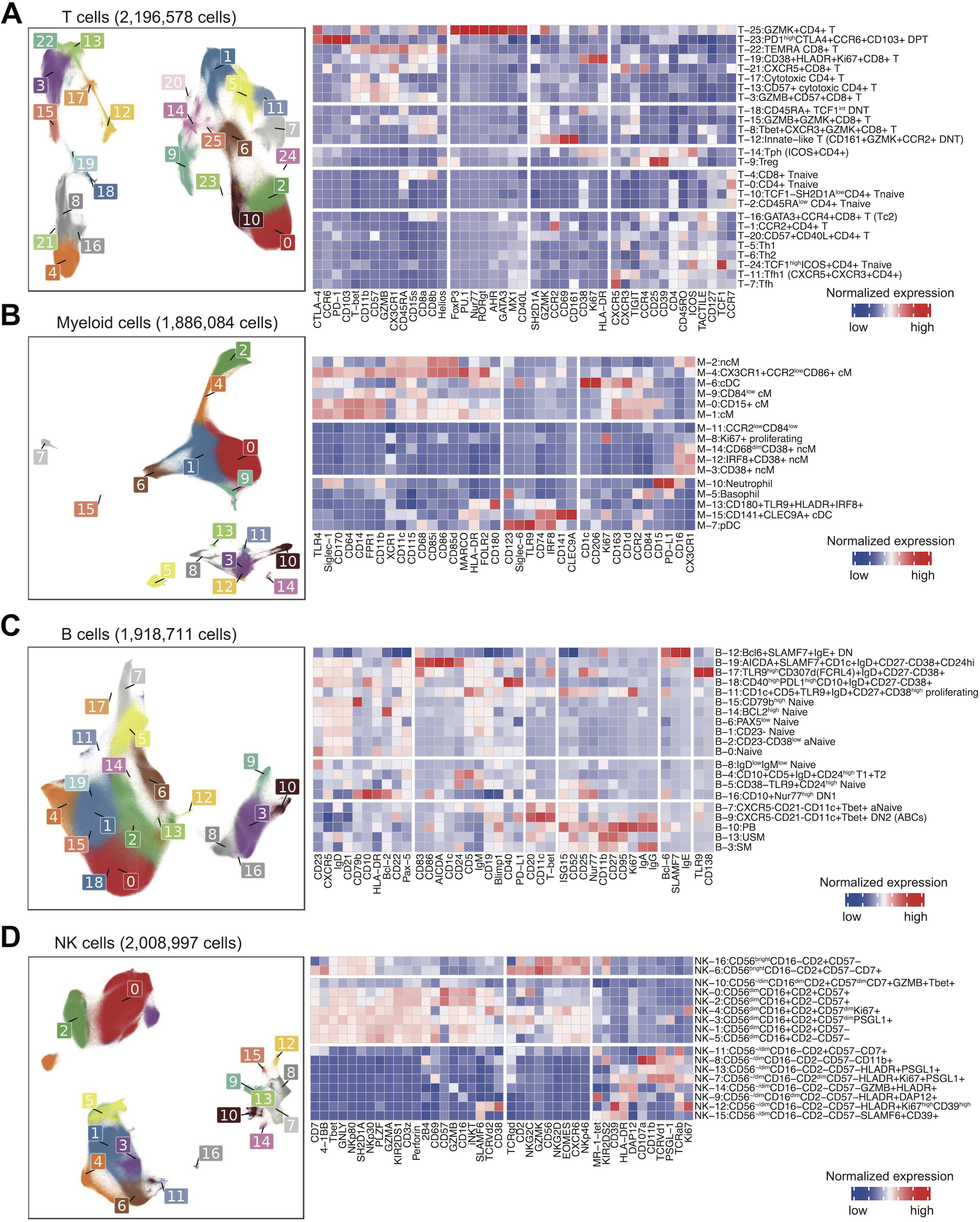
Cell-type-specific clustering analysis revealed 79 distinct cell states. **A-D.** Cell-type-specific immune proteomic reference colored by fine-grained cell states in the UMAP space. For each cell type, the heatmap shows the average expression distributions of key variable proteins in each cluster across samples, scaled within each cell cluster. Clusters are ordered by protein expression pattern using hierarchical clustering.

Among myeloid cells, we observed classical monocytes (M-0, M-1, M-4, M-9) and nonclassical monocytes (M-2), as well as conventional dendritic cell populations (M6, M15), and a separate plasmacytoid dendritic cell clusters (M-7) (**Fig.2B**, **Extended Data Fig.4**). In addition, we observed a small population of neutrophils (M-10), likely representing low density granulocytes, as well as a small population of basophils (M-5). B cells predominantly segregated into IgD+ naive (B-0-2, B-5-8, B-14, and B-15), and CD27+ memory B populations (B-3, B-13), while distinct populations of CXCR5^low^CD21^low^CD11c+ activated naïve B cells (B-7) and CD11c+ age-associated B cells (ABCs) (DN2) (B-9) clustered separately (**Fig.2C**, **Extended Data Fig.5**). NK cells separated predominantly into CD56^dim^ and CD56^bright^ populations (**Fig.2D**, **Extended Data Fig.6**). In all, many of the disease-relevant immune cell phenotypes are detectable in the peripheral blood from the At-Risk RA individuals.

### Unique immunophenotypes characterize At-Risk RA subpopulations

To systematically quantify cell abundances and their differences between At-Risk individuals and controls, we identified expanded cell neighborhoods within At-Risk individuals (n = 52) compared with healthy controls (n = 48) accounting for technical batches and demographic variables, including clinical site, age, and sex (**Methods**). This integrative approach not only gains power but also further evaluates the specific At-Risk status findings in a more generalized manner. We used this approach to identify specific immune phenotypes that are altered in At-Risk individuals compared to controls in T, myeloid, B, and NK cells, respectively as follows.

Within T cells, we observed skewed cell type abundances between At-Risk and control (*p* = 0.006) (**Fig.3A-B**, **Supplementary Table 4-5**). Specifically, cell neighborhoods among CCR2+CD4+ T cells (T-1), Th1 (T-5), Tfh1 (T-11), Tph (T-14), and a CXCR5+CD8+ T (T-21) were expanded in At-Risk, while both CD4+ and CD8+ naive T cells (T-0, T-2, and T-4) were depleted. CXCR5+CD8+ T (T-21) cluster was the most expanded cluster in At-Risk RA (OR = 3.33). This T cell population expressed high levels of chemokine receptors, but the cell frequency is relatively small (2,506 cells in blood of tested At-Risk individuals and controls) with high confidence intervals (2.11-5.28). Interestingly, the second most expanded At-Risk population was the CCR2+CD4+ T (T-1) cluster (OR = 1.47), a relatively large cluster (113,775 cells). In contrast, the GZMB+ effector CD8 T cell cluster (T-3) was depleted, as were three naive clusters (T0, T-2, T-4). Further, we investigated correlations between cell cluster abundances (**Extended Data Fig.7**) and found that the abundance of plasmablasts (PB, B-10) and Tph cells were significantly correlated (*R* = 0.42, *p* = 1e-9, **Fig.3C**), which further supports the hypothesis that Tph cells promote PB in inflammatory diseases^49, 50^. Intriguingly, the cell abundances of CCR2+CD4+ T and Tph cells were significantly correlated (*R* = 0.25, *p* = 4e-4, **Fig.3C**). To validate our findings, we analyzed an independent mass cytometry dataset for T cells generated from At-Risk RA (n=57) and controls (n=23) by mapping them to this original T cell reference to generate common cell state clusters; and further quantified the expansion and depletion of these T cell clusters in the validation dataset (**Fig.3D**, **Extended Data Fig.8**, **Supplementary Table 6-7, Methods**). Notably, we observed concordant expansion of these T cell clusters in At-Risk individuals between two independent cohorts, especially the significant ones expanded in At-Risk, including CCR2+CD4+ T, Tph, Th1, Tfh1, and CXCR5+CD8+ T cells (**Fig.3E**).

**Fig.3:**
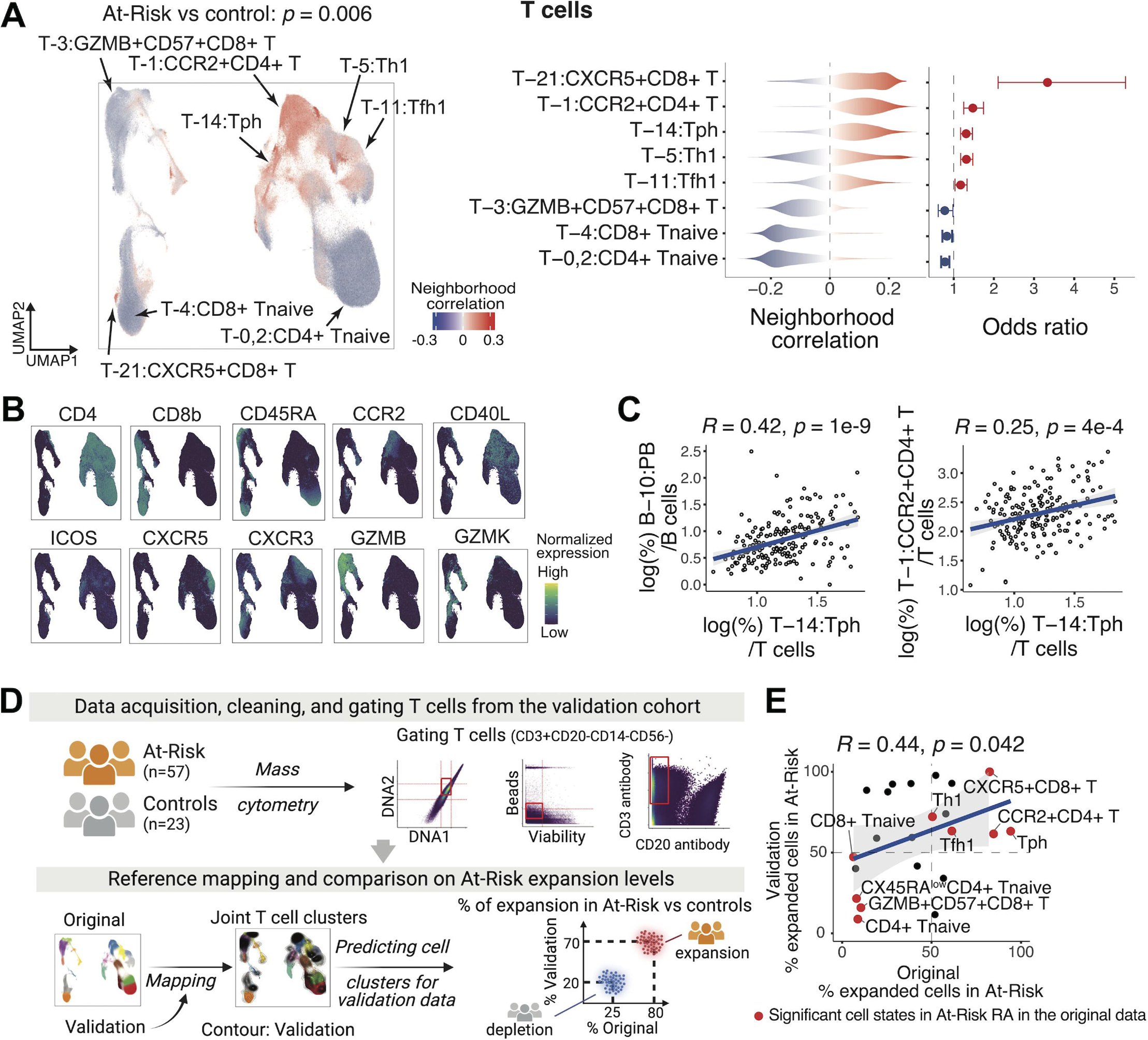
Identification of specific T cell populations that were associated with At-Risk. **A.** Associations of T cell neighborhoods with At-Risk vs controls (left). For all CNA-based association results, cells in UMAP are colored in red (expansion) or blue (depletion) and p-value is shown as well. Distributions of cell neighborhood correlations (center) and odds ratios (right) are shown. Error bars for odds ratio represent 95% confidence intervals, **B**. Expression of selected surface proteins within T cells are colored from dark-blue (low) to green (high), **C**. Scatter plot of cell type abundance correlations across individuals, **D.** Description of the validation dataset and analytical strategies, including reference mapping to the original T cell clusters, association test using CNA, and comparison of the proportion of expanded cells (neighborhood correlation > 0) in At-Risk (vs control) between two independent datasets by clusters. **E**. Scatter plot of the proportion of expanded cells in At-Risk by clusters, with x-axis for the original T cell panel and y-axis for the validation dataset. Red colored dots represent significant cell clusters in the original T cell panel. All statistical association tests are adjusted for age and sex. Correlation coefficients and p-values were obtained from Spearman’s correlation test.

To investigate signatures of CCR2+CD4+ T cells, we reanalyzed external single-cell RNA-seq (scRNA-seq) data obtained from human blood^51^, and identified differentially expressed genes (DEGs) (adjusted *p* < 0.05) in CCR2+ compared with CCR2-within CD4+ effector memory T cells (**Extended Data Fig.9**). All 338 DEGs were significantly upregulated in CCR2+CD4+ T cells, including *IFNγ* and *IL17RE* (**Supplementary Table 8**). Moreover, we reanalyzed a single-cell multimodal dataset from 70 RA synovial biopsies^37^. We found that *CCR2* was expressed on multiple T cell populations that infiltrating RA synovium, including Tph cells, CD4+CD161+ memory T cells, and CD4+IL17R+CCR5+ T cells, consistent with the idea that CCR2 may help induce immune cell migration in inflamed synovium^52^ (**Extended Data Fig. 10**).

Furthermore, we observed other immune cell phenotypes altered in At-Risk individuals, including expansion of CD15+ cM (M-0) (OR = 1.30, *p* = 0.001) in myeloid cells (**Fig.4A-B**), and expansion of PAX5^low^ naive B cells (B-6) (OR = 1.35, *p* = 0.009) uniquely in the FDR+ACPA+ At-Risk subgroup, the highest risk group for RA (**Fig.4C-D**). To investigate the phenotype of PAX5^low^ naive B cells, we compared the expression of activation markers that decrease with B cell activation (CD21 and CD23)^53, 54^ in the largest naive cluster (B-0). Notably, PAX5^low^ naive B cells were characterized by lower expression of CD21 and CD23 than conventional naive B cells, suggesting an activated state of PAX5^low^ naive B cells (**Fig.4E**). The PAX5^low^ naive B cluster (B-6) also expressed higher IgM protein expression levels than conventional naive B cells (B-0) (**Fig.2B**, **Extended Data Fig.5**). The activated naïve B cells that expanded in SLE were characterized by low expression of surface CXCR5, CD21, and CD23, in addition to high expression of CD11c and T-bet^55^, suggesting this might be a different subpopulation compared with the one expanded in SLE. Within NK cell clusters, we observed an expansion of the CD56^dim^CD16+CD2+CD57^dim^ proliferating (Ki67+) cluster (NK-4), and a depletion of CD56^dim^CD16+CD2-CD57-cells (NK-5) (**Fig.4F-G**).

**Fig.4:**
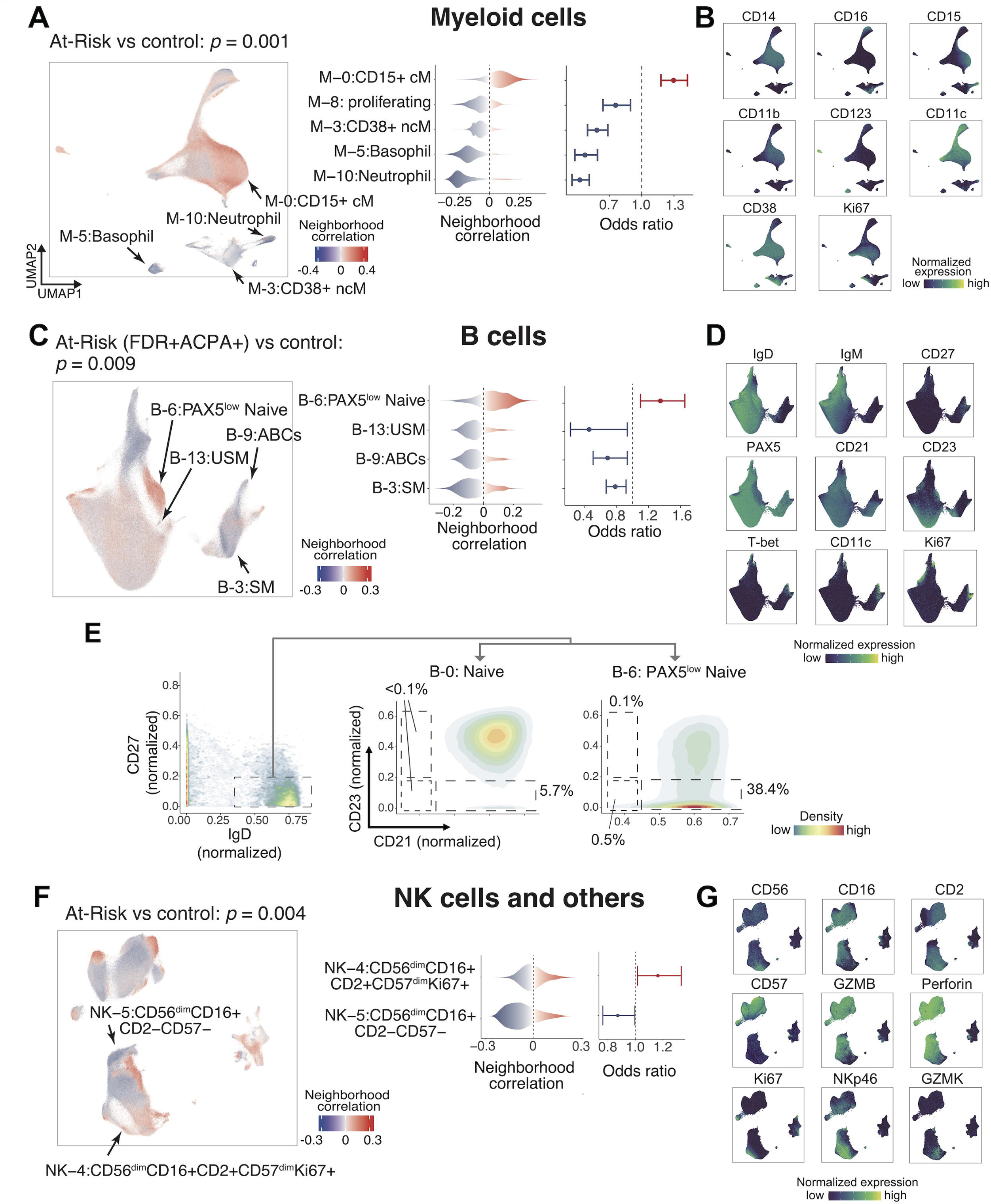
Identification of different myeloid, B cell and NK cell populations that were associated with At-Risk. **A.** Associations of myeloid cell neighborhoods with At-Risk vs controls (left), distributions of cell neighborhood correlations (center), and odds ratios (right) are shown, **B.** Expression of selected surface proteins within myeloid cells are colored from dark-blue (low) to green (high), **C.** Associations of B cell neighborhoods with At-Risk vs controls (left), distributions of cell neighborhood correlations (center), and odds ratios (right) are shown, **D**. Expression of selected surface proteins within B cells, **E**. Distributions of activation markers (CD21 and CD23) antibody staining in the conventional naive B cell cluster (B-0) and the PAX5^low^ naive B cell cluster (B-6). Low expression of both CD21 and/or CD23 indicates activated B cells, **F**. Associations of NK cell neighborhoods with At-Risk vs controls (left), distributions of cell neighborhood correlations (center), and odds ratios (right) are shown, **G**. Expression of selected surface proteins within NK cells. All the statistical association tests are adjusted for age and sex. For all CNA-based association results, cells in UMAP are colored in red (expansion) or blue (depletion) and p-value is shown as well. Error bars for odds ratio represent 95% confidence intervals.

### ACPA status is a main driver of the cellular heterogeneity in At-Risk individuals

Both the presence of serum ACPA and a family history of RA confer risk of RA^2^; however, it is unclear whether these two risk factors are associated with similar or distinct cellular changes in the pre-RA stage. Thus, we systematically characterized enrichment levels of 79 immune cell states in the ACPA+ At-Risk including FDR+ and FDR-, ACPA-FDR+ At-Risk, ACPA+ RA, and ACPA-RA compared with controls, respectively. Consequently, we identified 13 cell states that exhibited either significant enrichment (OR > 1) or depletion (OR < 1) in these individual groups (adjusted *p* < 0.05), suggesting different immune signatures according to ACPA status (**Fig.5**, **Supplementary Table 4**). Importantly, Tph cells (T-14), a population expanded in RA synovium, were found to be expanded in blood from both ACPA+ and ACPA-At-Risk subgroups, while CCR2+CD4+ T cells (T-1) were expanded specifically in the ACPA+ At-Risk subgroup. In addition, the CXCR5+CD8+ T cells (T-21) were expanded in both ACPA+ and ACPA-At-Risk individuals while not enriched in RA patients. This may suggest a pre-RA-specific immunophenotype, which warrants validation with longitudinal cohorts. In contrast, CD15+ cM (M-0) was expanded only in ACPA-At-Risk, while CD15-cM (M-1) was expanded only in ACPA+ At-Risk. Further, we observed a trend of expansion of ABCs (B-9) and PB (B-10) only in ACPA+ RA, while PAX5^low^ naive (B-6) was expanded in ACPA+ At-Risk, highlighting potential phenotypic differences between ACPA+ At-Risk and RA (**Extended Data Fig.11**). Through sensitivity analysis, we found that the FDR-ACPA+ At-Risk associations are concordant when comparing site matched controls and controls from multiple clinical sites (*R* = 0.52, *p* = 2e-4) (**Extended Fig. 12**). Similarly, the FDR+ACPA+ At-Risk associations were significantly concordant (*R* = 0.29, *p* = 0.033) (**Extended Fig. 12**). In all, these analyses suggest that these overabundant immune phenotypes (e.g., CCR2+CD4+ T cells) unique for the ACPA+ At-Risk subpopulation are maintained across multiple types of analysis.

**Fig.5:**
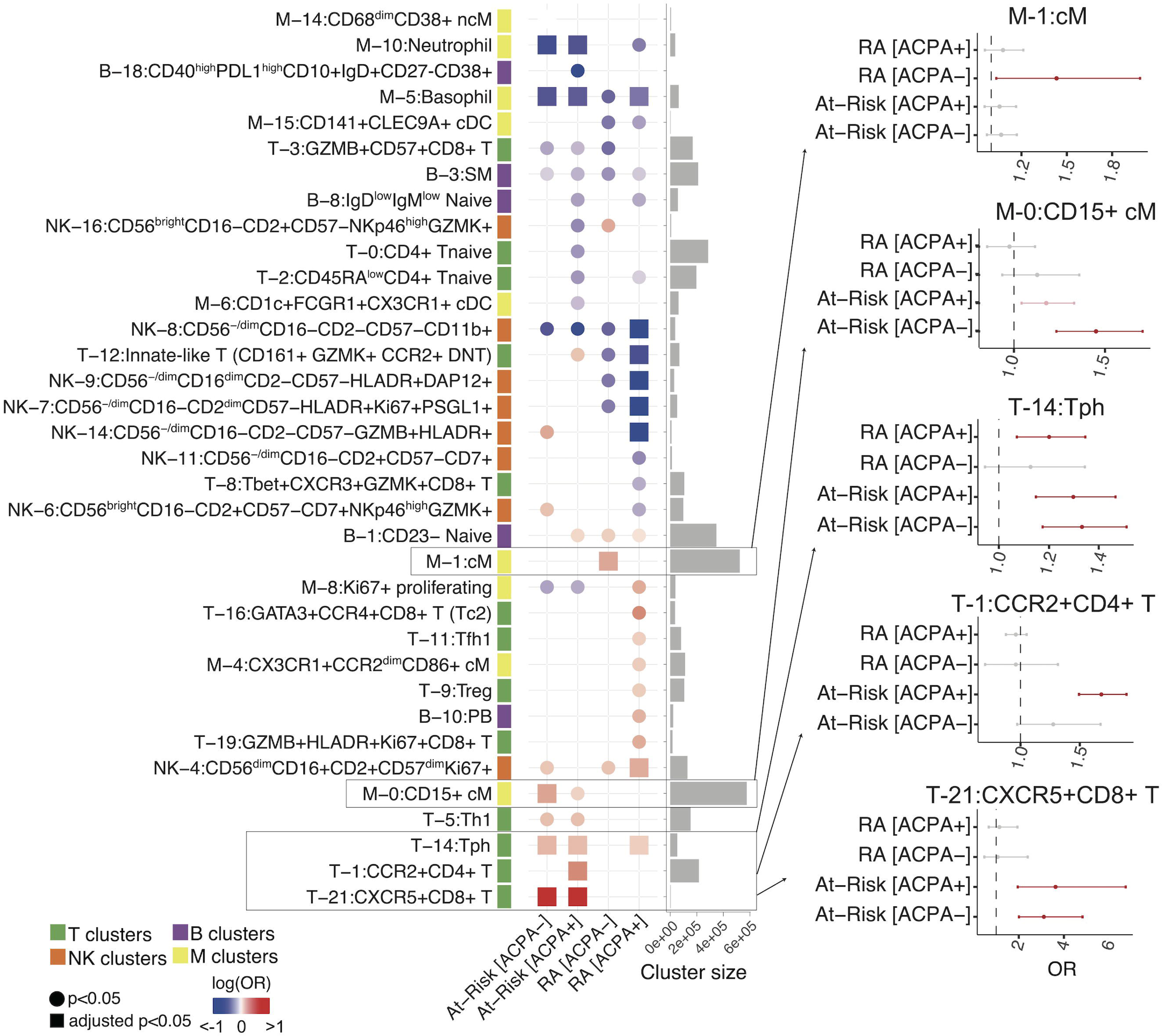
ACPA-status specific analysis reveals unique populations for different disease statuses. Heatmap shows association with each subgroup upon ACPA status in At-Risk and RA (vs controls) for each cell type. Only clusters with *p* < 0.05 are shown. Circles represent *p* < 0.05 and squares represent adjusted *p* < 0.05. Adjusted p-values were calculated by the Benjamini and Hochberg method. Cell types are colored in red (expanded) or blue (depleted). Error bars on selected cell populations represent 95% confidence intervals. All the results in this analysis are adjusted for age and sex.

### An RA immunophenotype score model to quantitatively classify disease statuses

To date, family history and ACPA as well as rheumatoid factor (RF) titers have been commonly utilized in clinical practice to assess the risk of developing RA in At-Risk populations^56–58^. With the development of high-throughput sequencing technologies, large-scale genome-wide association studies have provided a strategy for quantifying risk by summing the disease susceptibility alleles that individuals have^59^. However, these methods do not incorporate pathological immunophenotypes or measure their contribution to future risk for a single individual level.

To address this need, we developed the “RA immunophenotype score” classification method and evaluated its performance in our large-scale and cross-sectional single-cell proteomic data. We formulated the RA immunophenotype score by multiplying coefficients (𝛽), representing the degree of “RA-relevant” quantitatively obtained from the comparison of baseline RA at initial visit (vs controls), by corresponding cell state abundances (𝑋) (**Fig.6A**, **Methods**). First, to ensure the validity of applying this formula to each of the major immune cell types and further aggregating them into the RA immunophenotype model, we investigated whether cell type-specific cell neighborhoods are skewed between RA and controls using co-varying neighborhood analysis and ensuring the significance (all *p* < 1e-3). Then, to select RA-relevant cell states and define coefficients for each fine-grained cluster, we calculated the odds ratio as the enrichment score accounting for age, sex, and site, and filtered by adjusted *p* < 0.05 (**Methods**). In all, we identified 10 fine-grained cell clusters which are significant and incorporated into the RA immunophenotype model (**Fig.6B**). Lastly, we multiplied the generated enrichment score by cell type abundance for each individual, and summed them across all significant clusters by individuals. Note that our RA immunophenotype score model was trained on individuals from established RA and healthy controls. Thus, we further evaluated the RA immunophenotype score model regarding its performance in distinguishing At-Risk individuals, which were not used for modeling RA immunophenotype score, from healthy controls (**Fig.6C**). We demonstrated that the RA immunophenotype score can significantly separate At-Risk individuals from controls (AUC = 0.646; 95% CI: 0.534-0.758) (**Fig.6D**). Comparing altered cell clusters in At-Risk and RA, Tph cells were expanded in both At-Risk and RA, while CCR2+CD4+ T and GZMB+HLADR+Ki67+CD8+ T cells were uniquely enriched in At-Risk RA only and RA only, respectively (**Fig.6E**). We expect to apply this score in longitudinal studies to determine whether it can improve the prediction of the future development of clinical RA.

**Fig.6:**
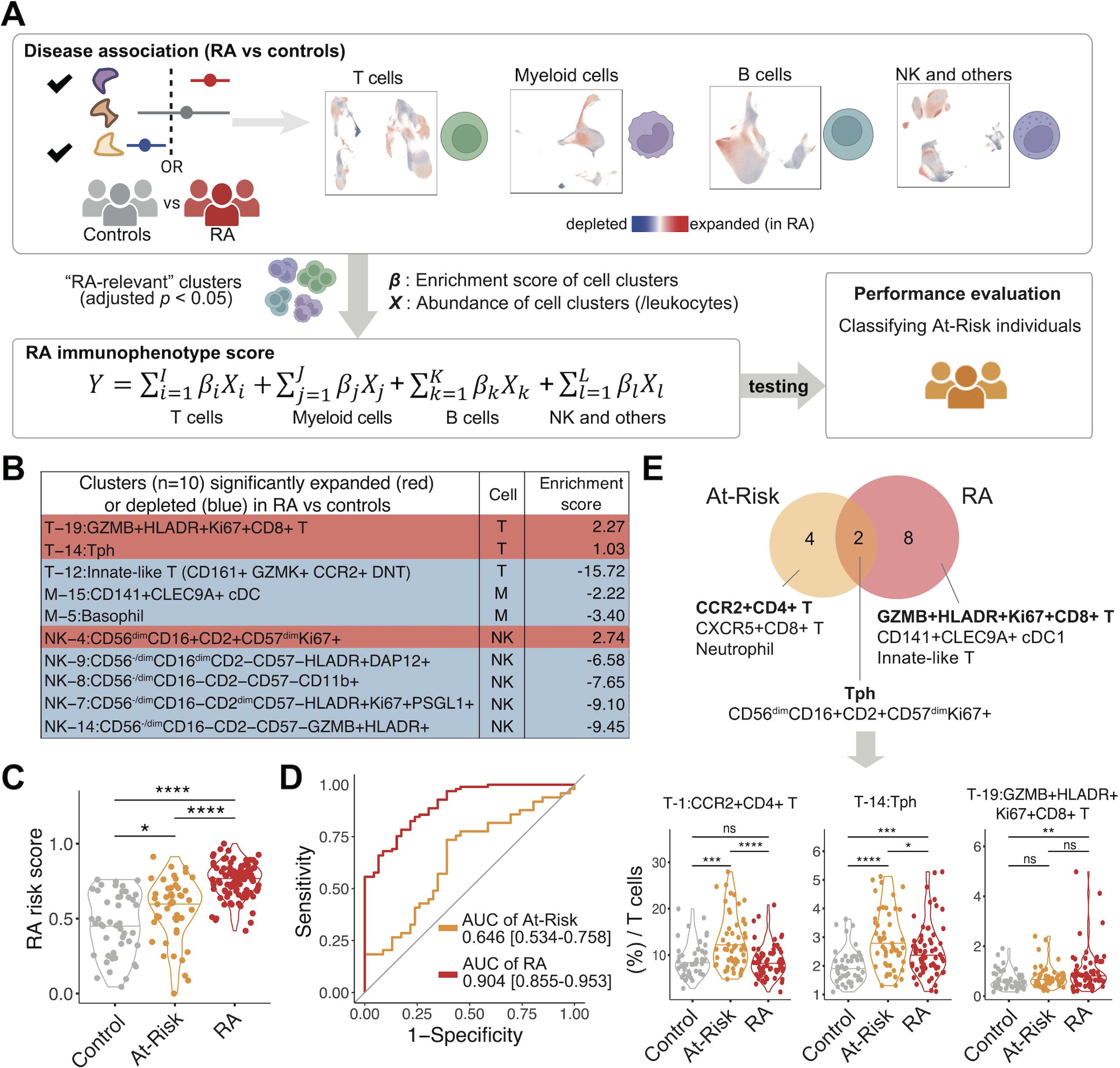
RA immunophenotype score utilizing RA-specific cell type abundances to quantify and distinguish At-Risk individuals from control. **A.** Pipeline for developing RA immunophenotype score. Associations of each RA-specific cell type neighborhood (vs control). For each cell type, all p-values from the CNA test were *p* < 1e-3. We incorporated clusters that are significantly associated with RA (adjusted *p* < 0.05) to model the RA immunophenotype score. Here, all the analyses are adjusted for age and sex. We calculated RA immunophenotype score based on cell type abundances multiplied by corresponding major cell type proportions and enrichment scores (Methods) for each cell type, **B.** Significant cell clusters that are either expanded (enrichment score greater than 0) or depleted (enrichment score less than 0) in RA vs controls and their enrichment scores. These are the clusters incorporated in the RA immunophenotype score model, **C.** Distribution of RA immunophenotype score across individual samples from RA, At-Risk, and controls; **** *p* < 0.0001, * *p* < 0.05, **D.** Receiver operating characteristic (ROC) analysis to evaluate the classification performance of RA immunophenotype score in distinguishing At-Risk from control. Areas under the curve (AUC) with 95% confidence intervals were described, **E.** Summarized Venn diagram of significant (adjusted *p* < 0.05) cells expanded or depleted in At-Risk and RA, respectively; **** *p* < 0.0001, *** *p* < 0.001, ** *p* < 0.01, * *p* < 0.05, ns non-significant.

## Discussion

We constructed an At-Risk landscape of immune cell atlas in blood utilizing comprehensive surface proteins of large-scale single-cell proteomics (> 8,000,000 cells). Through robust computational modeling and integrative analyses, we discovered cell clusters from different immune cell compartments that are significantly altered in At-Risk individuals. Given the limited availability of pre-RA cohorts and the challenge to harmonize cross-site single-cell data, our integrative and disease association strategies can be easily generalized to maximize the power to address similar disease progression questions. We further developed a disease classification model, designated the RA immunophenotype score, designed to quantify the RA-relevant immune signatures at the individual level, an approach which is currently needed to improve disease prediction biomarkers and further accelerate preventive strategy development.

During the past decade, research on established RA has been focusing on genetics and transcriptional regulation, as well as gene and protein expression, which have uncovered biologically meaningful signatures^33, 37, 59–65^. Single-cell transcriptomic and multimodal analysis have revealed high granular cell populations in the inflamed synovium to pinpoint phenotypes that characterize tissue inflammation^33, 37^. Recent clinical trial studies that examine synovial heterogeneity using bulk RNA-seq suggest that treatment response may depend on the specific immune composition in the tissue^66, 67^. More importantly, translating already identified RA-relevant signatures from tissue to inform prognosis or even predict clinical RA onset in the clinic is still challenging due to the knowledge gap of immunophenotypes in the At-Risk stage. Our study is among the first unbiased reports using mass cytometry to characterize the immune heterogeneity within different subsets of At-Risk populations. We detected already-known pathogenic cell populations and also uncovered novel phenotypes. For example, we identified and further defined the expansion of Th1 and Tfh1 in At-Risk individuals, which is consistent with the predominance of Th1 response over Th2 in RA^68, 69^. Although Tph cells are known to be increased in RA, especially in seropositive RA^44^, we reported here for the first time that these cells are also overabundant in the circulation of At-Risk individuals, including both ACPA+ and ACPA-At-Risk individuals. Given that the ACPA-At-Risk in our data are FDR and the ACPA status can reflect the genetic background in RA^70, 71^, the ACPA− At-Risk could include the population that will become ACPA+ in the future, and the expansion of Tph cells in this population may reflect this potential. Alternatively, the presence of expanded Tph cells may reflect either the genetic or environmental influences of being an FDR of an RA patient, regardless of ACPA status.

We also found an expanded CCR2+CD4+ T cluster, and our analysis using an external single-cell dataset further demonstrated that these CCR2+CD4+ T cells in the blood could be IFNγ-producing CD4+ T cells with effector functions like Th17 phenotypes (IL17RE+), intriguing their involvement in mucosal origin of RA endotype^21^. Multiple large-scale bulk RNA-seq datasets also support that *CCR2* can be expressed by Th17 cellse^72–74^. *CCR2*-expressing Th17 cells are known to produce inflammatory cytokines, including IL-17 and IFNγ, in the central nervous system leading to autoimmune encephalomyelitis in model mice^75^. On the other hand, since *CCR2* is expressed not only in Th17 but also in Th1, Th2, and Treg^72–74^, detailed study on CCR2+CD4+ T cells to define their fine-scaled phenotypes is necessary in the future.

We found that the CXCR5+CD8+ T cell population, expressing multiple chemokine receptors (CXCR3, CX3CR1, and CCR4), was expanded in At-Risk. Previous research shows that the CXCR5-expressing follicular CD8+ T cells migrate into B-cell follicles and are important in the response to chronic viral infection^76–83^ and cancers^84–86^. This cell population in our dataset is relatively small, thus further experimental validation is needed to investigate whether they are consistent with previously reported CXCR5-expressing CD8+ T cells. Assessment of the T cell receptor repertoire of this T cell population in the pre-RA phase using the approach in our parallel study^87^ may also enhance our understanding of their origin and function.

A few studies have investigated the efficacy of DMARDs in the pre-RA period^19, 20, 88, 89^. One critical challenge is how to precisely identify the high-risk individuals that actually need intervention, because only a subset of At-Risk individuals will develop RA in the near term, and there are not yet well-established strategies for prevention^19, 20, 88, 89^. Further, because preventive studies are aimed at people who have not yet developed RA, a balance must be struck between expected adverse events and efficacy, and the use of predictive markers for treatment response may be useful^90^. For example, the B cell-directed therapy on preclinical individuals delays RA onset^20^. Here, we identified a PAX5^low^ naive B cell phenotype overabundant in FDR+ACPA+ At-Risk. This B cell population may be a novel pre-plasmablast state given that PAX5 is a key transcription factor in B cell development but is repressed during plasma cell differentiation^91, 92^. In parallel, the clinical trial for the T cell costimulatory molecule inhibitor CTLA4-Ig in pre-RA is ongoing^88^. The CCR2+CD4+ T cells we identified and the Tph cells have the potential of being considered as predictive markers given their expansion in ACPA+ At-Risk and highly expressed proteins for B cell help. We also found a myeloid population, CD15+ classical monocytes, particularly expanded in seronegative At-Risk individuals. It is not known whether any of these At-Risk individuals will develop RA with seroconversion in the future, thus further investigating the mechanisms of these phenotypes in RA history may help delineate a more precise predictive marker.

Advances in computational integration algorithms facilitated the cross-institution, cross-tissue, and cross-disease analyses, revealing underlying shared mechanisms and pathways in immune-mediated diseases^34, 36, 42, 93–96^. Most of these studies primarily integrated data on mRNA expression levels and chromatin states. Here we presented single-cell proteomics references that comprehensively incorporated 119 individuals from At-Risk and established RA together with 57 At-Risk RA individuals as validation, which can serve as references to query immune phenotypes involved during RA progression and conversion (e.g., early or established RA, before or post-treatment). Our study also includes samples at different time points from the same RA individuals, although they are not sufficient to test the changes of unique cell populations caused by treatments in this study (**Supplementary Table 1**). Further, our references will help clarify the blood-tissue comparison to elucidate the circulation pathways of the pathogenic immune phenotypes identified in this study and their migration mechanisms between blood and tissue.

Based on these intriguing results, we developed a novel RA immunophenotype score that can measure phenotypical changes of key immunophenotypes to successfully distinguish At-Risk, established, and healthy controls. Note that a caveat is that for optimal effect, such scoring systems must also incorporate the years of follow-up to determine which specific At-Risk individuals actually develop RA. Longitudinally acquired samples to determine the rate and timing of disease conversion were not obtained for this study. Hence, it is necessary for our RA immunophenotype score prediction model and the identified immunophenotypes to be further evaluated in other large longitudinal preclinical cohorts. In addition, the next stages of study should focus on determining how the phenotypes described herein relate to other pre-RA phenotypes, including alterations in the lung mucosa^26^ and gut microbiome^22^, among others.

## Methods

### Subject recruitment and clinical data collection

The Accelerating Medicines Partnership (AMP) Network for RA and SLE constructed a cross-sectional cohort. PBMC samples from RA and controls were collected from eleven clinical sites across the United States and two sites in the United Kingdom. RA individuals were diagnosed based on the 2010 ACR/EULAR criteria^97^. Healthy controls from three clinical sites were tested to be ACPA and RF and they were negative. The original samples from At-Risk individuals without inflammatory arthritis seen in rheumatology clinics were collected from the University of Colorado Anschutz Medical Campus. The collection occurred over the course of a 45-month period from October 2016 to February 2020. The study was performed in accordance with protocols approved by the institutional review board. Demographics, clinical data, and measurements of laboratory data were performed at the baseline visit. Data collected include age, sex, race, RF or ACPA status, RA treatments, and tender and swollen joint counts. For At-Risk individuals, we defined ACPA-positive as anti-CCP3 and/or anti-CCP3.1 titer >= 20 units. For established RA, we defined ACPA-positive as anti-CCP1, anti-CCP2, and/or anti-CCP3 titer more than upper limit of normal range of each clinical site. ESR and CRP were measured for established RA patients using commercial assays in each institution’s clinical laboratory. Disease activity for each RA patient was calculated using a CDAI^98^. For validation, samples from At-Risk individuals and controls were collected from the same two clinical sites, including the University of Colorado Anschutz Medical Campus and Brigham and Women’s Hospital; and the At-Risk individuals from both sites were defined and characterized using the same classification strategies, family history and the positivity of ACPA, for validation.

### Collection of samples and processing

Samples were shipped to the central AMP RA/SLE Biorepository, Oklahoma Medical Research Foundation Biorepository, until sample collection was complete. They were then transited to the central pipeline site at the Brigham and Women’s Hospital CyTOF Antibody Resource and Core, where samples were thawed and processed in 23 batches.

### Mass cytometry antibody staining and quality control

All PBMC samples from 167 individuals (established RA (n=67), At-Risk RA (n=52) and controls (n=48)) were thawed in a 37 °C water bath for 3 minutes and then mixed with 37 °C thawing media containing: RPMI Medium 1640 (Life Technologies #11875-085) supplemented with 5% heat-inactivated fetal bovine serum (Life Technologies #16000044), 1 mM GlutaMAX (Life Technologies #35050079), antibiotic-antimycotic (Life Technologies #15240062), 2 mM MEM non-essential amino acids (Life Technologies #11140050), 10 mM HEPES (Life Technologies #15630080), 2.5 x 10^-5^ M 2-mercaptoethanol (Sigma-Aldrich #M3148), 20 units/mL sodium heparin (Sigma-Aldrich #H3393), and 25 units/mL benzonase nuclease (Sigma-Aldrich #E1014). 100 μL aliquots of each sample post-thaw were mixed with PBS (Life Technologies #10010023) at a 1:1 ratio to be counted by flow cytometry. Between 0.5 – 1.0 x 10^6^ cells were used for each sample. All samples were transferred to a polypropylene plate (Corning #3365) to be stained at room temperature for the rest of the experiment.

The samples were spun down and aspirated. Rhodium viability staining reagent (Standard BioTools #201103B) was diluted at 1:1000 and added for five minutes. 16% stock paraformaldehyde (Fisher Scientific #O4042-500) was diluted to 0.4% in PBS and added to the samples for five minutes. After centrifugation and aspiration, Human TruStain FcX Fc receptor blocking reagent (BioLegend #422302) was used at a 1:100 dilution in cell staining buffer (CSB) (PBS with 2.5 g bovine serum albumin [Sigma Aldrich #A3059] and 100 mg of sodium azide [Sigma Aldrich #71289]) for 10 minutes followed by incubation with conjugated surface antibodies (each marker was used at a 1:100 dilution in CSB, unless stated otherwise) for 30 minutes. All antibodies were prepared and validated by the Harvard Medical Area CyTOF Antibody Resource and Core (Boston, MA).

After centrifugation, samples were resuspended with culture media. 16% stock paraformaldehyde (Fisher Scientific #O4042-500) dissolved in PBS was used at a final concentration of 4% for 10 minutes to fix the samples before permeabilization with the FoxP3/Transcription Factor Staining Buffer Set (ThermoFisher Scientific #00-5523-00). The samples were incubated with SCN-EDTA coupled palladium barcoding reagents for 15 minutes followed by incubation with Heparin (Sigma-Aldrich #H3149-100KU) diluted 1:10 in PBS. Samples were combined and filtered in a polypropylene tube fitted with a 40µm filter cap. Conjugated intracellular antibodies were added into each tube and incubated for 30 minutes. Cells were then fixed with 4% paraformaldehyde for 10 minutes.

To identify single cell events, DNA was labeled for 20 minutes with an 18.75 μM iridium intercalator solution (Standard BioTools #201192B). Samples were subsequently washed and reconstituted in Cell Acquisition Solution (CAS) (Standard BioTools #201240) in the presence of EQ Four Element Calibration beads (Standard BioTools #201078) at a final concentration of 1×10^6^ cells/mL. Samples were acquired on a Helios CyTOF Mass Cytometer (Standard BioTools). The raw FCS files were normalized to reduce signal deviation between samples over the course of multi-day batch acquisitions, utilizing the bead standard normalization method established by Fink et al^99^. The normalized files were then compensated with a panel specific spillover matrix to subtract cross-contaminating signals, utilizing the CyTOF based compensation method established by Chevrier et al^100^. These compensated files were then deconvoluted into individual sample files using a single cell based debarcoding algorithm established by Zunder et al^101^. Pre-analysis of CyTOF staining data included a Gaussian gating strategy^102^, gating on singlet cells by residual versus DNA staining, gating on bead-negative cell events, and gating on all live cells (Rhodium-negative).

### Downsampling cells for all mononuclear cells, T, and myeloid panels

T cells and myeloid cells consist of a large proportion of peripheral blood. In order to save time and computational resources for downstream analysis without missing important cell states, we downsampled cells by randomly selecting cells according to individuals for analyses for all mononuclear cells, T cells, and myeloid cells as follows;

1. If 10% of total cells > 10,000, we will keep 10% of total cells
2. If 10,000 > 10% of total cells, we will keep 10,000 cells
3. If 10,000 > total cells, we will keep total cells without downsampling

For sensitivity analysis, we performed consistent clustering analysis and obtained biological cell clusters according to the proportions of downsampling (0.1%, 1%, 10%, 20%, 30%, 40%, 50%, 60%, 70%, 80%, 90%, and 100%).

### Protein expression normalization and dimensionality reduction

To minimize the effect of background on the measured signal, we normalized expression data by ArcSinh transformation of data using the cytofAsinh function in cytofkit R package with cofactor = 5 for each cell type. For dimensionality reduction, we then used truncated principal component analysis (PCA) as implemented in the prcomp_irlba function from the irlba R package and calculated 20 principal components (PCs) based on the normalized mass cytometry data. During PCA, we used the most highly variable proteins by removing 10% lowest variable proteins among cells because they are uninformative. We further corrected batch effects and heterogeneity within samples simultaneously with the HarmonyMatrix function from the harmony R package. We next projected the cells into two dimensions with UMAP^103, 104^ with default parameters.

### Graph-based clustering, differential protein expression, and cell type annotation

After batch correction, we constructed shared nearest neighbor graphs derived from the top 20 PCs and applied graph-based Louvain clustering^105^ at various resolution levels (0.3, 0.5, 0.7, 1.0). We selected optimized resolution values for each cell type (0.7 for T cells, 0.3 for NK cells, 0.3 for myeloid cells, 0.5 for B cells) based on silhouette width and manual check of expression of key proteins in each cluster to gain the biological interpretations that made the most sense. Unreliable clusters less than 30 cells in total were removed. In the end, we identified 26 T cell clusters (2,196,578 cells, 163 individuals), 20 B cell clusters (1,918,711 cells, 167 individuals), 17 NK clusters (2,008,997 cells, 160 individuals), 16 myeloid clusters (1,886,084 cells, 161 individuals), for a total of 79 clusters. We allocated cluster numbers based on cluster size. For each major cell type, we identified differentially expressed surface proteins by comparing cells from one cluster with all the other cells using wilcoxauc function in presto R package. We tested all proteins that were measured in each cell type. We present cluster-specific marker proteins and relative statistics in **Supplementary Table 3**. We then annotated identified clusters based on differentially expressed markers and relevant literature showing their biological functions in each cell type.

### Identification of cell populations that are significantly associated with specific clinical subgroups

We evaluated whether At-Risk or RA are associated with changes in the relative abundances of cell states within all mononuclear cells (coarse) and major cell type-specific manner (fine-grained). For each cell type, we applied multiple computational strategies, 1) cluster-based approach utilizing mixed effect model, Mixed-effects Association testing for Single Cells (MASC)^40^, and 2) cluster-free based approach which identifies dominant co-vary cell neighborhoods in cell type abundance across samples in one clinical group compared to the other, covarying neighborhood analysis (CNA)^41^. MASC is a statistical association strategy that uses single-cell logistic mixed-effect modeling to test individual cellular populations for their association by predicting the subset membership of each cell based on fixed effects and random effects. In MASC, a null model where the subset membership of every single cell is estimated by fixed and random effects without considering the case-control status of the samples was assumed. We then measured the improvement in model fit when a fixed effect term for the case-control status of the sample was included with a likelihood ratio test. This framework allowed us to evaluate the significance and effect size of the case-control association for each cluster while controlling for inter-individual and technical variability. In our analyses, we performed MASC using the MASC() R function as follows:

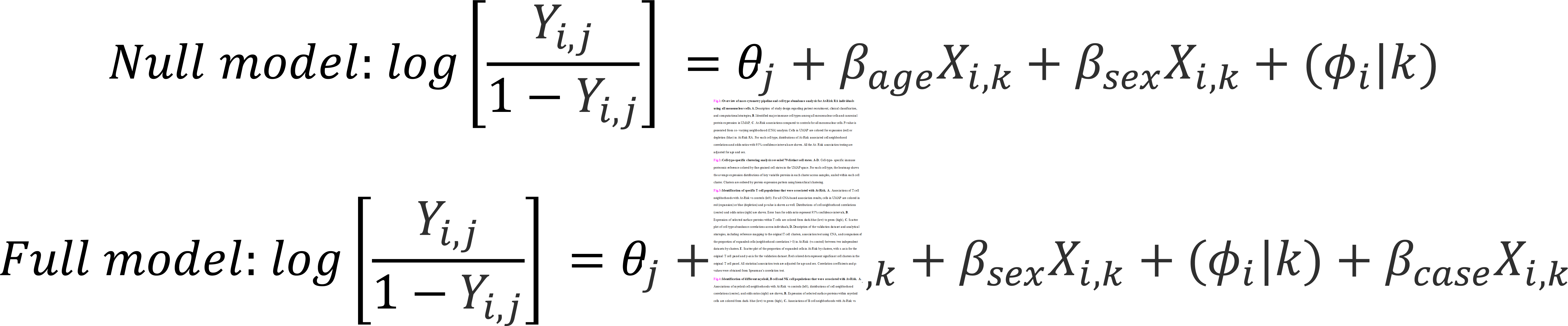

Here, *Y_i,j_* is the odds of cell *i* belonging to cluster *j* (major cell types for all mononuclear cells analysis and fine-grained cell types for each cell type analysis, respectively), *θ_j_* is the intercept for cluster *j*, *β_age_* and *β_sex_*indicate the fixed-effect of age and sex for cell *i* from k^th^ sample, respectively; (*ϕ_i_* |*k*) is the random effect for cell *i* from k^th^ sample, *β_case_* indicates the effect of k^th^ sample’s case-control status. We presented our results from MASC by odds ratio with an error bar indicating 95% confidence intervals for each cluster. The statistics are summarized in **Supplementary Table 4**.

It is noted that, for clusters with small cell numbers, statistics of MASC tend to have a wide range of confidence intervals and are unreliable, making it necessary to use the cluster-free method such as CNA^41^ in combination. We use CNA to define small cell neighborhoods in the batch-corrected harmonized low-dimensional embeddings and calculate that fractional abundance of cells from each sample in each neighborhood in a neighborhood abundance matrix (NAM). By decomposing the NAM with principal component analysis (PCA), CNA defines NAM-PCs within each cell type that capture axes of heterogeneity defined by groups of neighborhoods whose abundances vary in a coordinated manner. Next, we use CNA to perform two tests: associations between At-Risk vs control, and RA vs control, respectively. In practice, we used the association() function in the rcna R package with default parameters, while controlling for the “age” and “sex” as covariates. As CNA utilizes a permutation test, we obtained a significant association based on a global permutation *p* < 0.05. For visualization of local associations, we indicate the particular neighborhoods driving a global significant association. In the violin plots and UMAP plots, we colored neighborhood correlations, with red and blue indicating a positive and negative correlation, respectively. To highlight important cell neighborhoods from important cell states, we put transparent parameters according to the absolute value of correlation for each cell (from 0 [completely transparent] to 1 [no transparent]). The statistics of CNA results are in **Supplementary Table 5**.

### Reference mapping of independent mass cytometry T cells to the original T cell reference

We analyzed independent mass cytometry data obtained from blood of At-Risk (n=57), RA (n=20), and controls (n=23) enrolled from two clinical sites (University of Colorado and Brigham Women’s Hospital). Samples were shipped to the same central biorepository site until sample collection was complete. They were then transited to the central pipeline site, the same lab with the original sample processing, where samples were thawed and processed in 5 batches. After removing beads and dead cells by DNA gating, we gated T cells by CD3+CD20-CD56-CD14- and downsampled in the same way as the original T cell panel. To validate our findings in the original data, we then projected 1,022,630 T cells to the original T cell reference using the mapQuery() function based on 29 common proteins from the Symphony package. For reference building from the Harmony objects, we used the buildReferenceFromHarmonyObj() function. We predicted cell states for the query cells based on the 30 nearest cell neighbors using the knnPredict() function with k=30.

### Identifying gene signatures for the CCR2+CD4+ T cells

To infer the biological function of CCR2+CD4+ T cells in human blood, we explored gene signatures characterizing this population by reanalyzing publicly available single-cell RNA-seq dataset obtained from blood of healthy control. We downloaded the dataset (GSM3589419, resting cells from blood of donor A after negative selection to enrich CD3+ cells) from the Gene Expression Omnibus database with accession number GSE126030. To compare CCR2+CD4+ T cells and CCR2-CD4+ T cells, we defined them based on raw read count (CCR2+ effector CD4+ T cells, *CCR2* > 0 & *CD4* > 0 & *CD8A* = 0 & *CD8B* = 0 & *CCR7* <= 2; CCR2-effector CD4+ T cells, *CCR2* = 0 & *CD4* > 0 & *CD8A* = 0 & *CD8B* = 0 & *CCR7* <=2). We then normalized and scaled the data using NormalizeData function and ScaleData function from the Seurat R package. To identify differentially expressed genes, we used wilcoxauc function in presto R package. We defined differentially expressed genes if their adjusted *p* < 0.05. We present differentially expressed genes and relative statistics in **Supplementary Table 8**.

### Modeling RA immunophenotype score for each individual

To model RA immunophenotype score, we first ensured whether disease status (established RA or control) was associated with changes in the relative abundances of cell states within each of the four major cell types by applying CNA and confirmed all p-values for each cell type cluster that were smaller than 0.05. For established RA, we included all samples from individuals at baseline (n=66). Then, we applied MASC and defined “RA-relevant” cell clusters if their false discovery rate was smaller than 0.05. In total, 10 fine-grained cell clusters are selected to be incorporated into the RA immunophenotype model. To generate coefficients in the model for selected cell clusters, we developed the enrichment score by multiplying odds ratio from MASC by negative logarithmic-transformed p-value given that reliability of odds ratio depends on how robust they are. To generate RA immunophenotype score for each individual, we multiplied the abundance (among total mononuclear cells) of selected clusters by enrichment score for each cell type and summed them by individuals as follows;

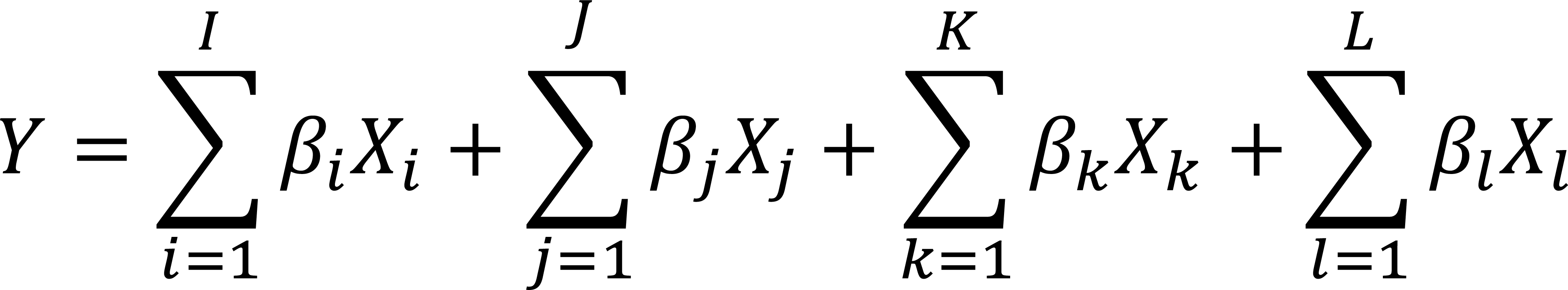

where *β* is enrichment score and *X* is cell cluster abundance among all mononuclear cells for each cell cluster (*i*, *j*, *k*, and *l* are cluster indexes for T, myeloid, NK, and B cells, respectively). For abundance, we performed logarithmic-transformation to minimize the effect from outliers. We further evaluated the performance of the RA immunophenotype score schema by applying on At-Risk individuals, which were not used for modeling RA immunophenotype score. We calculated and displayed the ROC (receiver operating characteristic curve) results regarding the capability of distinguishing At-Risk individuals from controls. The statistics of MASC used for modeling RA immunophenotype score for each cell type are in **Supplementary Table 4**.

## Acknowledgments

This work was supported by the Accelerating Medicines Partnership (AMP) in Rheumatoid Arthritis and Lupus Network. AMP is a public-private partnership (AbbVie Inc., Arthritis Foundation, Bristol-Myers Squibb Company, Foundation for the National Institutes of Health, GlaxoSmithKline, Janssen Research and Development, LLC, Lupus Foundation of America, Lupus Research Alliance, Merck Sharp & Dohme Corp., National Institute of Allergy and Infectious Diseases, National Institute of Arthritis and Musculoskeletal and Skin Diseases, Pfizer Inc., Rheumatology Research Foundation, Sanofi and Takeda Pharmaceuticals International, Inc.) created to develop new ways of identifying and validating promising biological targets for diagnostics and drug development. Funding was provided through grants from the National Institutes of Health (P30AR073750, UC2AR081032, UH2-AR067676, UH2-AR067677, UH2-AR067679, UH2-AR067681, UH2-AR067685, UH2-AR067688, UH2-AR067689, UH2-AR067690, UH2-AR067691, UH2-AR067694 and UM2-AR067678). Accelerating Medicines Partnership and AMP are registered service marks of the U.S. Department of Health and Human Services. This work was also supported by the Uehara Memorial Foundation Postdoctoral Fellowship, Grant-in-Aid for JSPS Overseas Research Fellows, and Mochida Memorial Foundation for Medical and Pharmaceutical Research (to J.I.); NIAMS R01 AR080659, NIAMS R01 AR077607, NIAMS P30 AR070253, and NIAMS P30 AR072577, Clinical Translational Science Award 1UL1TR002541-01 to Harvard University, and the Brigham and Women’s Hospital from the National Center for Research Resources (to J.A.S.); the PhRMA Foundation Faculty Starter Grant and the Arthritis National Research Foundation award (to F.Z.). We especially acknowledge people in the AMP RA/SLE Network: Arnon Arazi, Celine Berthier, Jill Buyon, Maria Dall’Era, Anne Davidson, Betty Diamond, Andrea Fava, Jennifer Grossman, Nir Hacohen, David Hildeman, Jeffrey Hodgin, Tiffany Hwang, Mariko Ishimori, Ken Kalunian, Diane Kamen, Matthias Kretzler, Holden Maecker, Rong Mao, Maureen McMahon, Fernanda Payan-Schober, Michelle Petri, Chaim Putterman, Daimon Simmons, Thomas Tuschl, David Wofsy, Steve Woodle, and Aaron Wyse.

## Author contributions

K.D.D, V.M.H., J.S., M.L.F., and J.M.N recruited patients, obtained samples, and curated clinical data for the SERA cohort study. J.A.J., M.B.B., S.R., W.A., L.B., V.B., S.G., L.D., G.S.F., H.P., J.M.B, L.B.H., D.T, A.F., C.P., J.H.A., L.M., and J.G. contributed to the procurement and processing of samples and design of the AMP RA/SLE study. K.D.D., V.M.H., and J.A.S. recruited and obtained samples for validation study. J.A.L., J.K., A.G., K.H., J.P., E.M., and B.H. designed and implemented the sample preparation, cell sorting, and mass cytometry data generation pipeline. J.I. led the computational and statistical analyses with support from T.G., A.G., and Y.C. For result interpretation, J.I., A.M.H., D.A.R., F.Z., and A.H.J. provided disease immunology inputs. J.I. and F.Z. wrote the initial draft. J.I., F.Z., D.A.R, K.D.D., and V.M.H. edited the manuscript. F.Z., D.A.R., J.A.L, K.D.D., and V.M.H. supervised the research. All authors participated in editing the final manuscript.

## Data availability

All raw and processed data will be available upon acceptance.

## Code availability

All source code to reproduce the analyses is available on GitHub (https://github.com/fanzhanglab/AtRiskRA_CyTOF/). Supplementary Information is available for this paper.

## Extended Data Figure legends

**Extended Data Fig.1: Downsampling strategy for large-scale mass cytometry dataset. A. O**ptimized downsampling schema developed for large-scale mass cytometry dataset to efficiently conduct downstream analysis without losing robustness, **B.** Sensitivity analysis for downsampling strategy. X-axis represents the proportion of downsampling cells. Y-axis represents the number of identified biologically meaningful cell clusters in each cell type using graph-based clustering.

**Extended Data Fig.2: Analytical pipeline applied to large-scale mass cytometry data. A.** Representative example of batch effect correction using myeloid panel. **B.** LISI scores in myeloid panel to measure mixture levels on race, clinical site, batch, and samples. After batch effect correction, the mixture level of clinical sites (median LISI = 4.45), technical batches (median LISI = 7.16) are significantly reduced compared to before correction (mean LISI = 4.31 for clinical sites, mean LISI = 5.53 for technical batchers (Wilcoxon test *p* < 0.01), **C.** Distribution of samples (top) and cell types (bottom) by batch, **D.** Analytical pipeline from expression data to cell embeddings in low-dimensional space using dimensionality reduction, **E.** Density plot using all mononuclear cells by batch. Cells from different batches but the same cell types are clustered together, **F.** Gating strategy for mass cytometry data to determine selected immune cell populations.

**Extended Data Fig.3: Expression of measured proteins in T cells panel.**

**Extended Data Fig.4: Expression of measured proteins in myeloid cells panel.**

**Extended Data Fig.5: Expression of measured proteins in B cells panel.**

**Extended Data Fig.6: Expression of measured proteins in NK cell panel.**

**Extended Data Fig.7: Correlation of abundance in blood between 79 cell types.** Correlation plot between cell type abundances. Cells are colored in red (positive) or green (negative) if their false discovery rate is less than 0.05.

**Extended Data Fig.8: Paired clusters after reference mapping using independent mass cytometry data for T cells.** We mapped T cells in the validation dataset onto the corresponding T cell reference from the original T cell panel to determine correspondent cell cluster annotations. **A.** Blue-red color scale in the heatmap indicates the log(OR) for a given pair of states (OR is the ratio of odds of mapping a cell cluster in the validation dataset to a given cluster of the original T cell panel compared to odds of mapping other cells in the validation dataset onto the same cluster of the original T cell panel), with higher values indicating greater correspondence. **B.** LISI scores of T cells from the validation data to measure mixture levels on clinical site, batch, and samples. After batch effect correction, the mixture level of clinical sites (median LISI = 1.85) and technical batches (median LISI = 3.56) are significantly increased compared to before correction (Wilcoxon test *p* < 0.01) suggesting the well mixture of cells in each T cell clusters, **C.** Cell count after assigning predicted cell clusters based on the original T cell panel, **D.** Average expression distributions of variable key proteins in each cluster across samples, scaled within each cell cluster.

**Extended Data Fig.9: Meta-analysis a single-cell RNA-seq blood dataset enriching CD3+ T cells to infer gene signatures of CCR2+CD4+ T cells. A.** Workflow and results focusing on CCR2+ vs CCR2-within CD4+ T cells using data from GSM3589419 in GSE126030, B. UMAP colored by fine-grained cell states in the single-cell RNA -sequencing data, C. Expression of selected genes of interest.

**Extended Data Fig.10: Expression of *CCR2* mRNA in the synovium of RA patients**. **A.** T cell clusters identified in the synovium of RA patients^37^ in the UMAP space. The annotations of *CCR2*-expressing clusters are labeled. **B.** *CCR2*-expressing cells in the UMAP. Expressing cells are colored in blue.

**Extended Data Fig.11:** Association with each clinical subgroup according to ACPA status in At-Risk and RA (vs controls) for selected B cell populations. Error bars represent 95% confidence intervals for the odds ratio. Statistical results are obtained by adjusting for age and sex.

**Extended Data Fig.12: Sensitivity analysis for different control groups from multi-clinical sites. A.** Density plot by family history and ACPA status according to cell types, **B.** Correlation plot of odds ratios comparing At-Risk RA subgroups with FDR−ACPA− controls (y-axis, n=8) from the SERA cohort (y-axis) or healthy controls from other clinical sites (x-axis, n=40). Dots are colored by immune cell types. Of total association tests, 77 cell clusters were included; outliers of the odds ratio (top 99%ile and bottom 1%ile) or size of clusters are lower than 25%ile among all clusters, and results with infinite confidence intervals for the odds ratio were excluded. Statistical results are adjusted for age and sex. Correlation coefficients and p-values were obtained by Spearman’s correlation test.

## Supplementary Tables

**Supplementary Table 1.** Characteristics of enrolled controls, At-Risk, RA and different At-Risk subpopulations.

**Supplementary Table 2**. Mass cytometry panels for each immune cell type.

**Supplementary Table 3**. Differentially expressed proteins in cell types.

**Supplementary Table 4**. Statistics of cluster-based associations with At-Risk subgroups or RA (vs controls).

**Supplementary Table 5**. Statistics of single-cell neighborhood associations with At-Risk subgroups or RA (vs controls).

**Supplementary Table 6**. Mass cytometry panel for T cells in the validation dataset.

**Supplementary Table 7**. Characteristics of enrolled subjects in the validation dataset.

**Supplementary Table 8**. Differentially expressed genes in CCR2+CD4+ vs CCR2-CD4+ using single-cell RNA-seq.

