## Extended Data Figures for "Deep immunophenotyping reveals circulating activated lymphocytes in individuals at risk for rheumatoid arthritis"

**A**

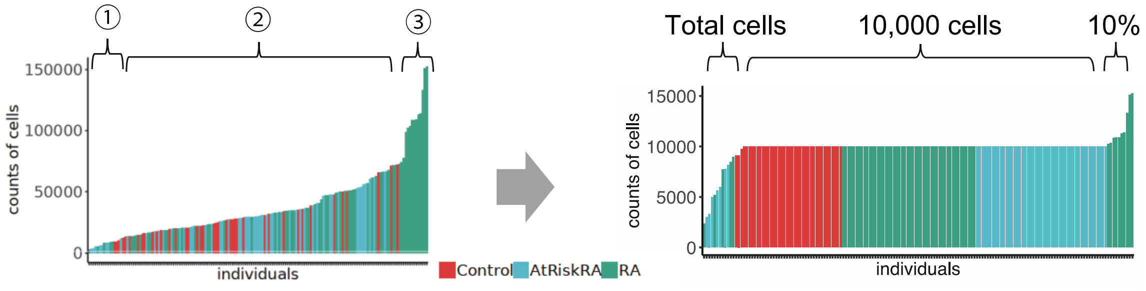


T and Myeloid panels: randomly selecting cells according to individuals as following;
① if 10% of total cells > 10,000 → 10% of total cells are kept
② if 10,000 > 10% of total cells → 10,000 cells are kept
③ if 10,000 > total cells → total cells are kept (without downsampling)

**B**

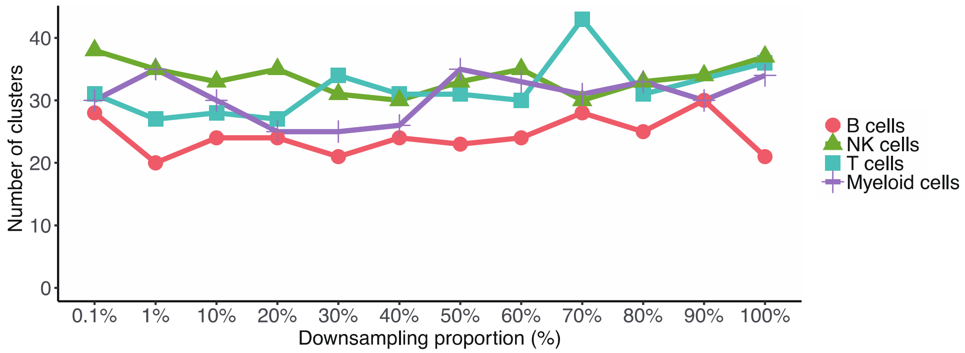


**Extended Data Fig.1: Downsampling strategy for large-scale mass cytometry dataset. A**. Optimized downsampling schema developed for large-scale mass cytometry dataset to efficiently conduct downstream analysis without losing robustness, **B**. Sensitivity analysis for downsampling strategy. X-axis represents the proportion of downsampling cells. Y-axis represents the number of identified biologically meaningful cell clusters in each cell type using graph-based clustering.

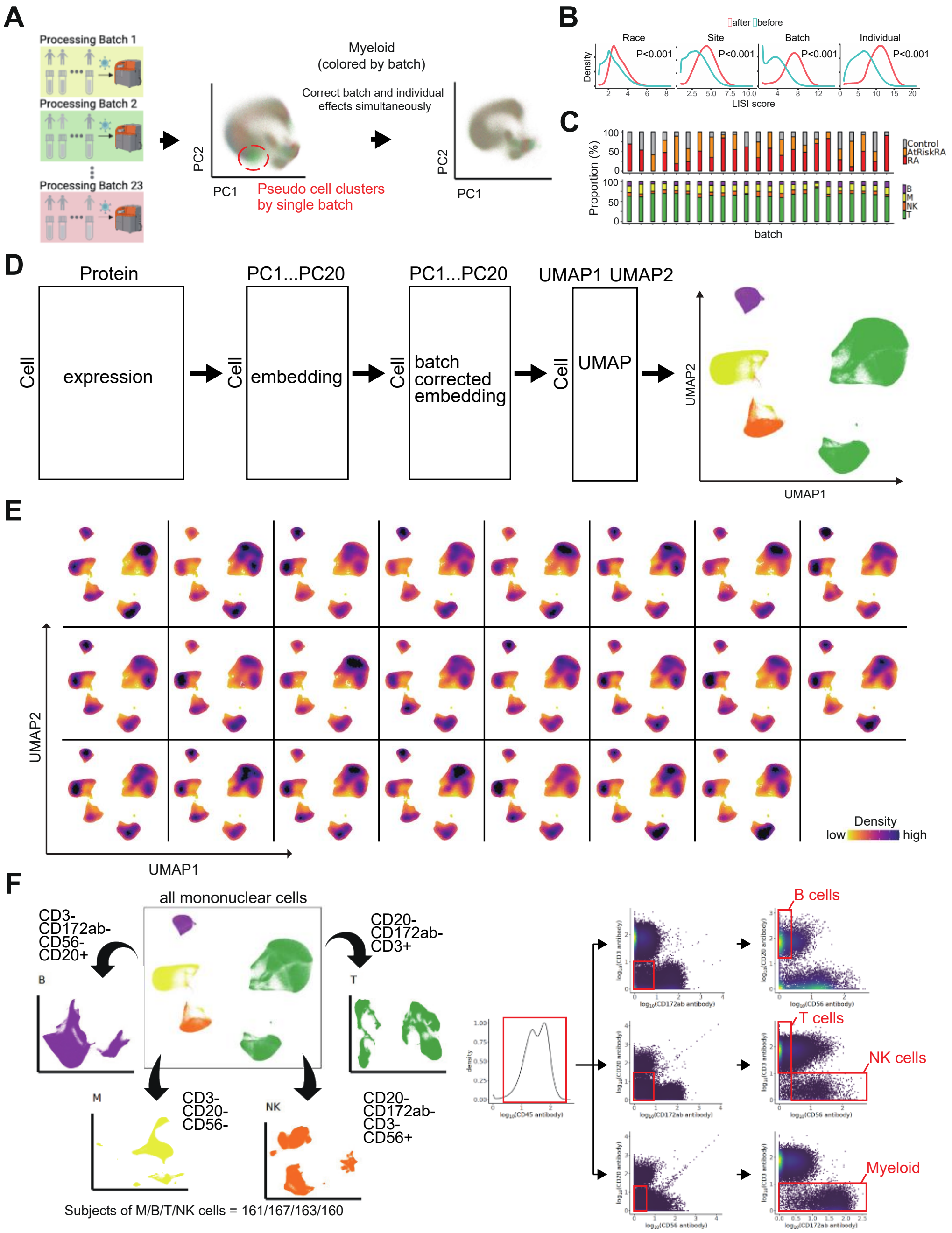


**Extended Data Fig.2: Analytical pipeline applied to large-scale mass cytometry data. A**. Representative example of batch effect correction using myeloid panel. **B.** LISI scores in myeloid panel to measure mixture levels on race, clinical site, batch, and samples. After batch effect correction, the mixture level of clinical sites (median LISI = 4.45), technical batches (median LISI = 7.16) are significantly reduced compared to before correction (mean LISI = 4.31 for clinical sites, mean LISI = 5.53 for technical batchers (Wilcoxon test $p < 0.01$), **C**. Distribution of samples (top) and cell types (bottom) by batch, **D**. Analytical pipeline from expression data to cell embeddings in low-dimensional space using dimensionality reduction, **E**. Density plot using all mononuclear cells by batch. Cells from different batches but the same cell types are clustered together, **F**. Gating strategy for mass cytometry data to determine selected immune cell populations.

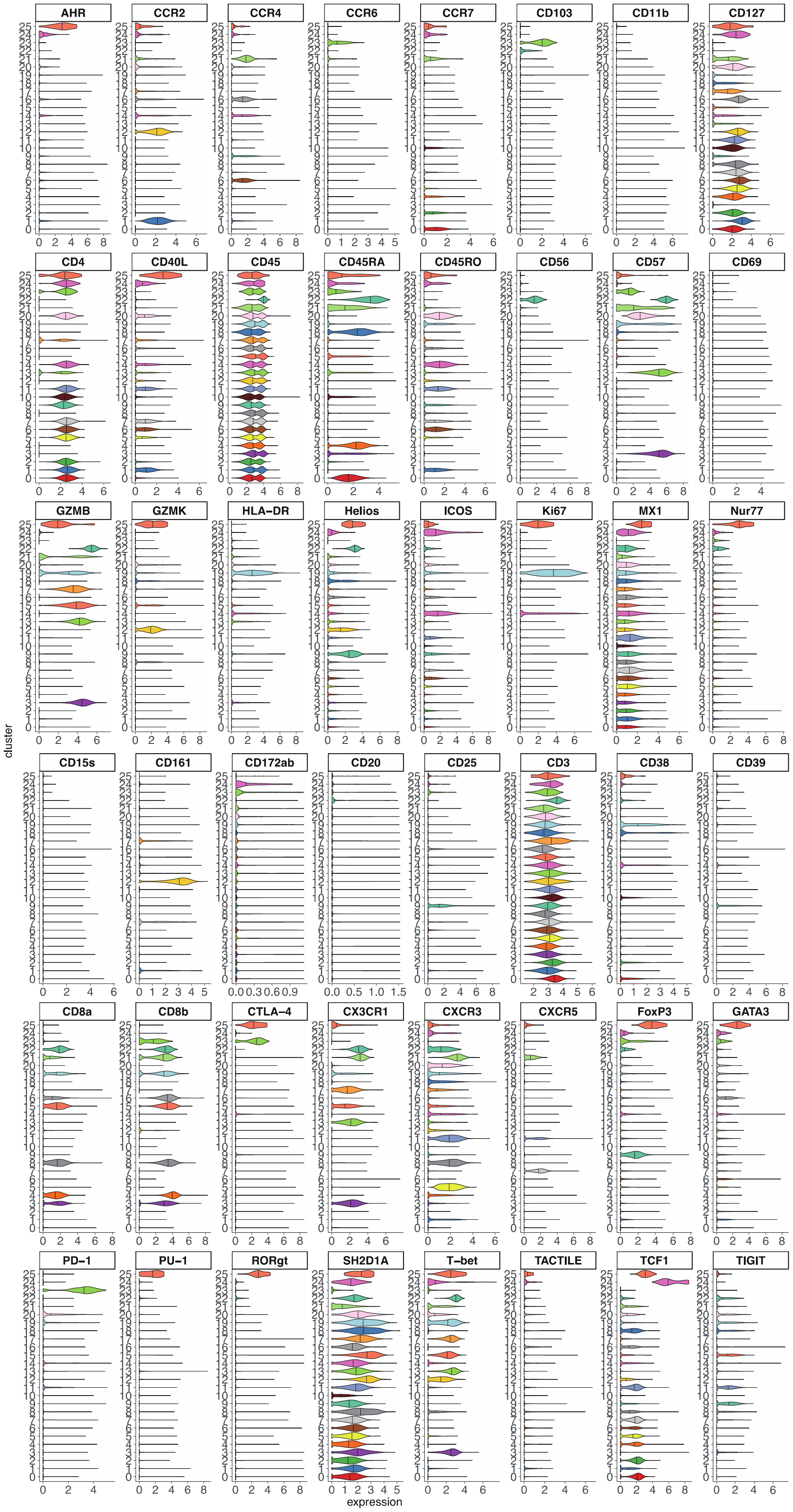


**Extended Data Fig.3: Expression of measured proteins in T cells panel.**

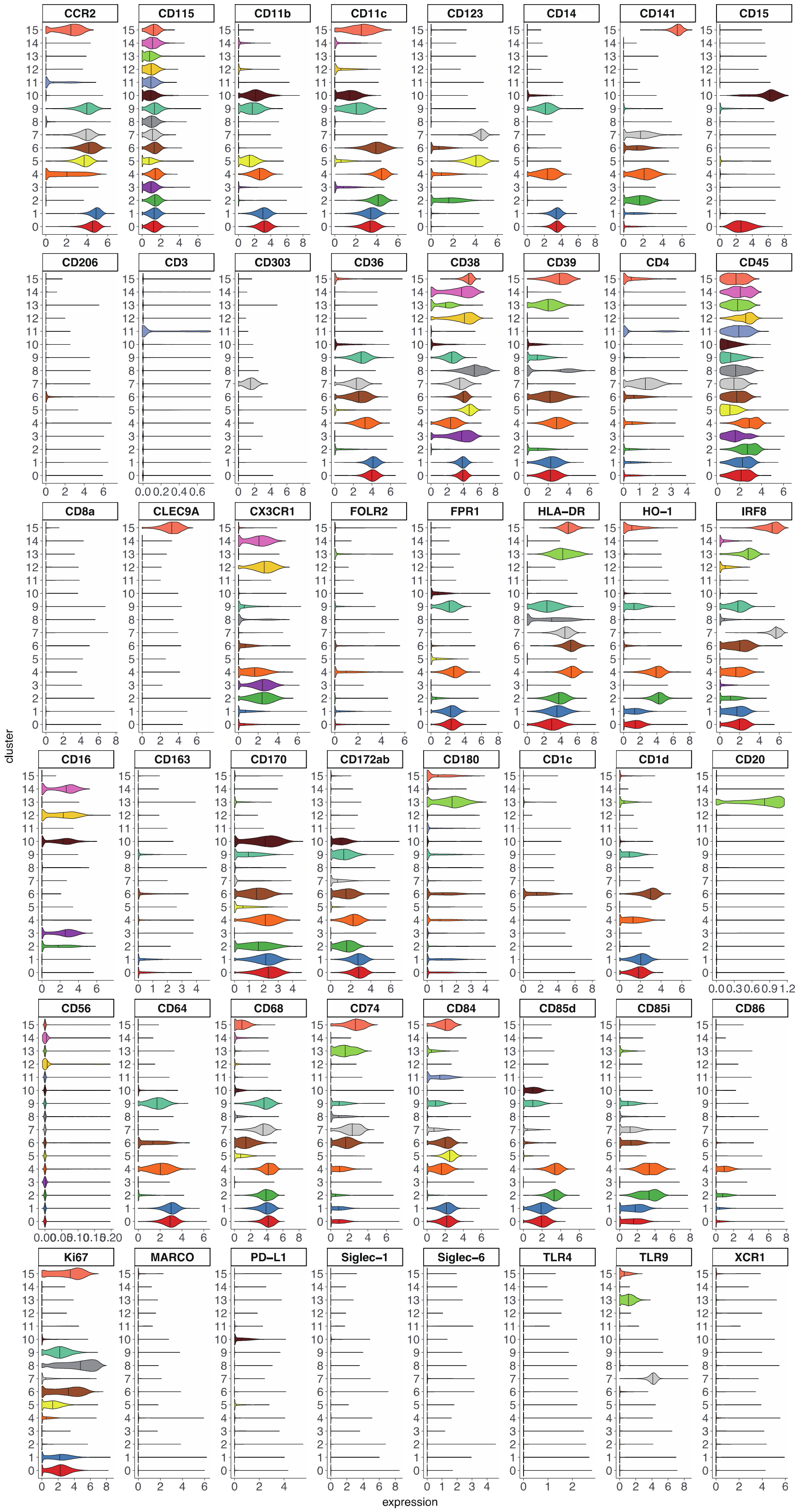


**Extended Data Fig.4: Expression of measured proteins in myeloid cells panel.**

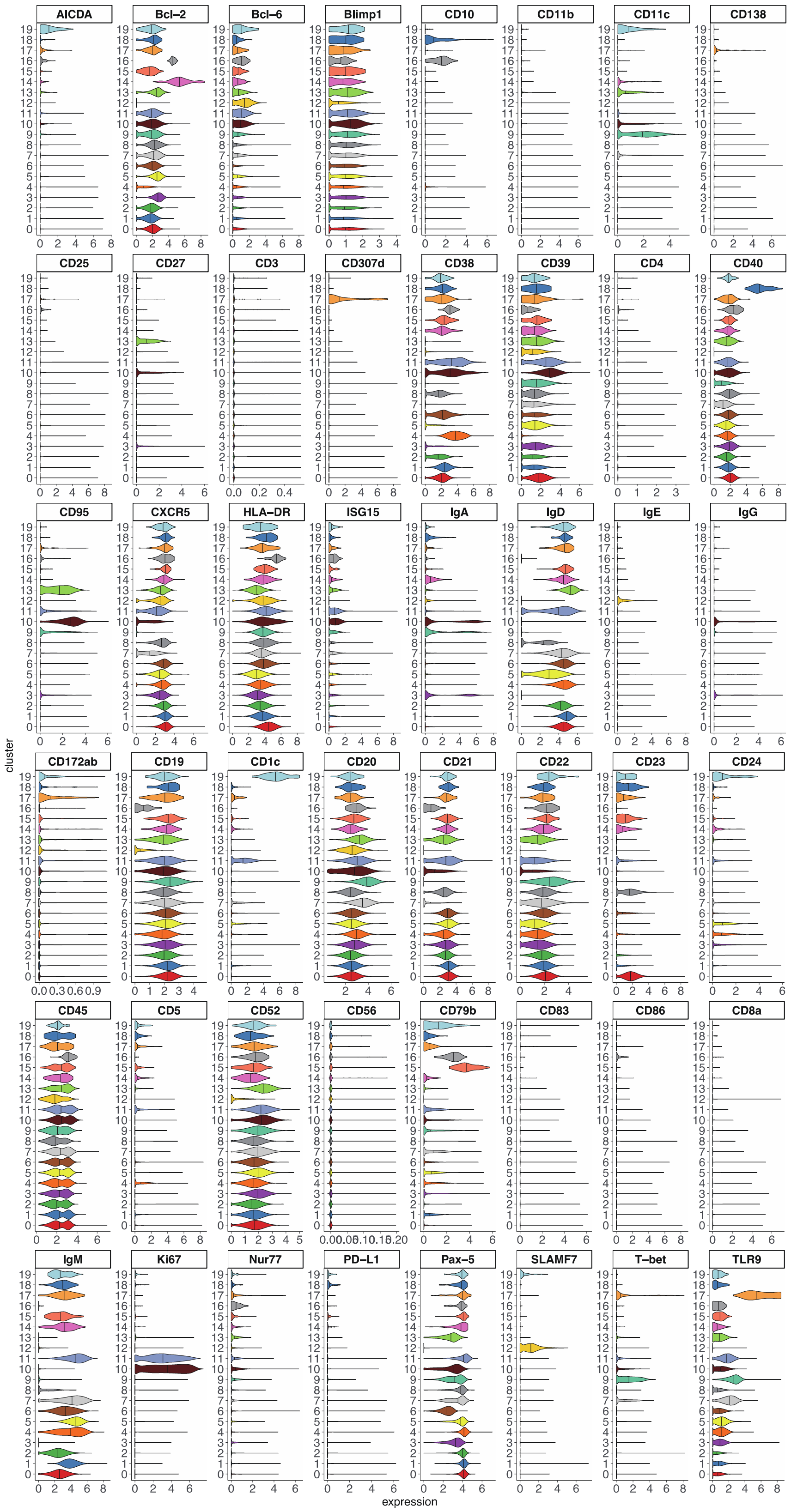


**Extended Data Fig.5: Expression of measured proteins in B cells panel.**

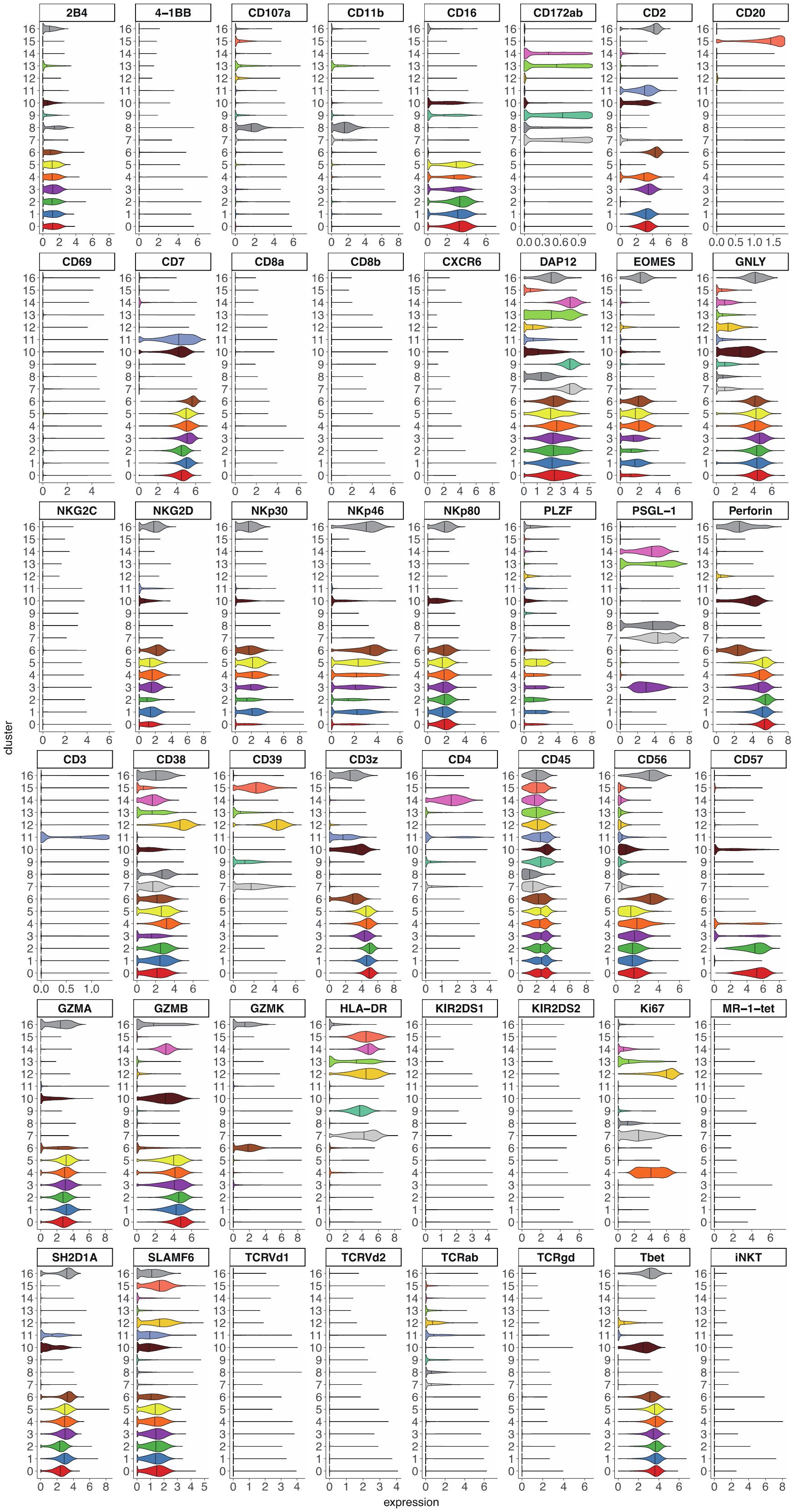


**Extended Data Fig.6: Expression of measured proteins in NK cell panel.**

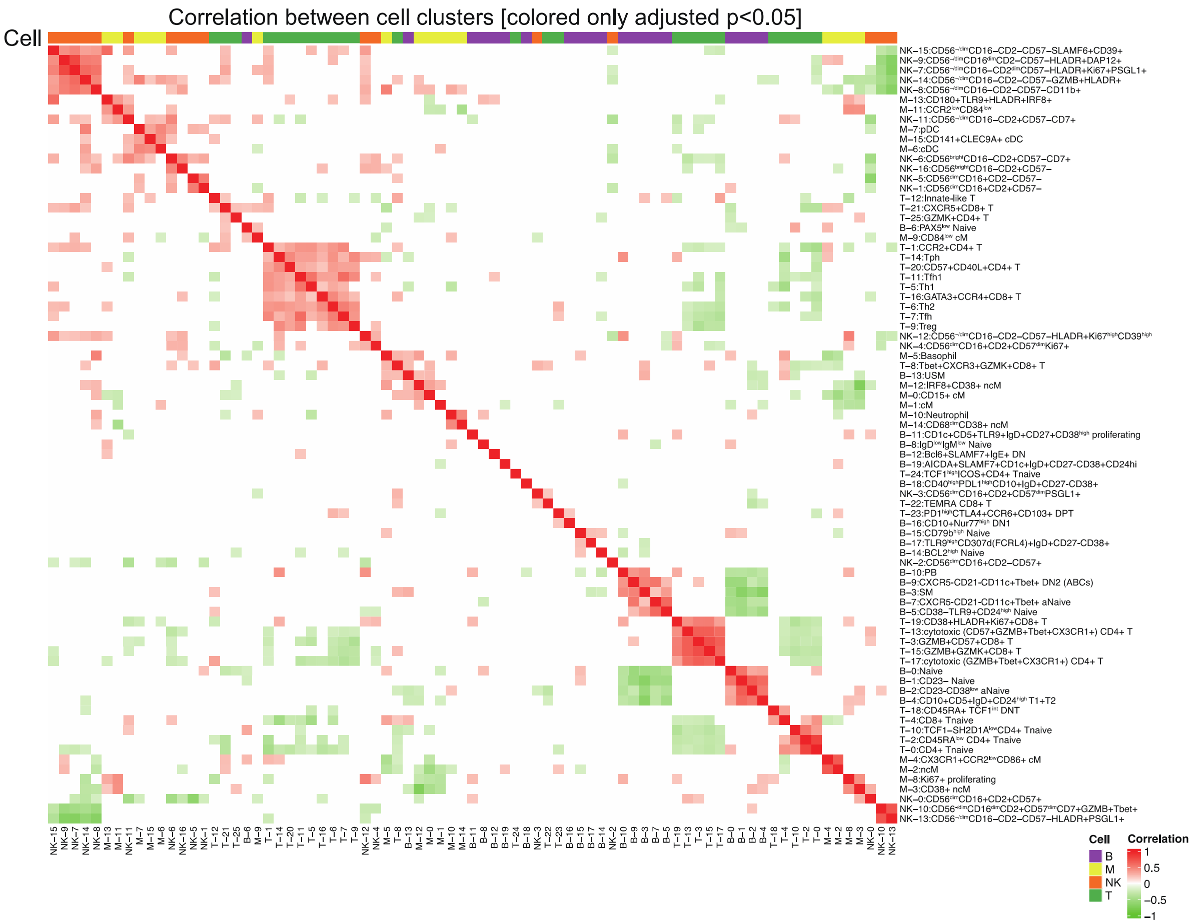


**Extended Data Fig.7: Correlation of abundance in blood between 79 cell types.** Correlation plot between cell type abundances. Cells are colored in red (positive) or green (negative) if their false discovery rate is less than 0.05.

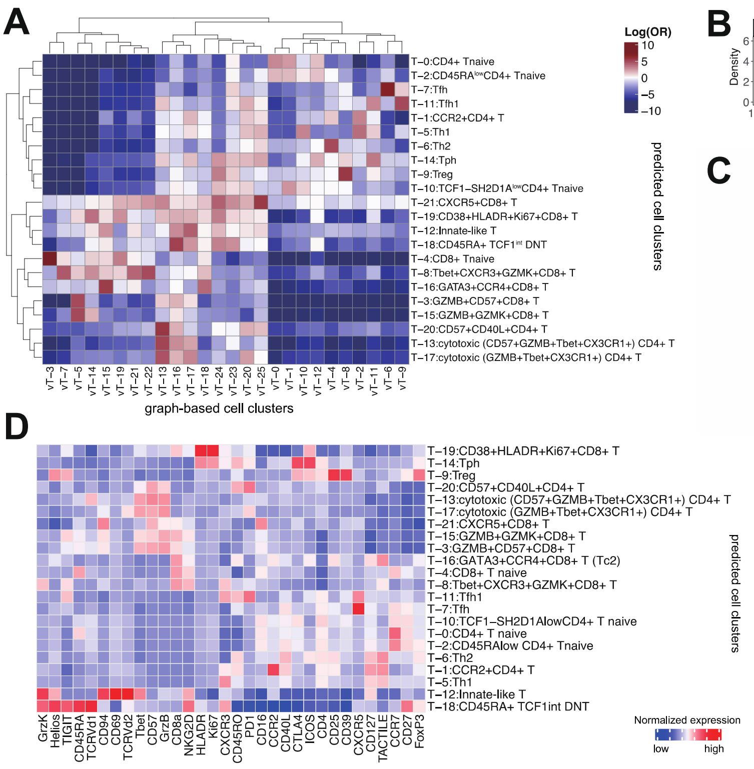


**Extended Data Fig.8: Paired clusters after reference mapping using independent mass cytometry data for T cells.** We mapped T cells in the validation dataset onto the corresponding T cell reference from the original T cell panel to determine correspondent cell cluster annotations. **A.** Blue-red color scale in the heatmap indicates the log(OR) for a given pair of states (OR is the ratio of odds of mapping a cell cluster in the validation dataset to a given cluster of the original T cell panel compared to odds of mapping other cells in the validation dataset onto the same cluster of the original T cell panel), with higher values indicating greater correspondence. **B.** LISI scores of T cells from the validation data to measure mixture levels on clinical site, batch, and samples. After batch effect correction, the mixture level of clinical sites (median LISI = 1.85) and technical batches (median LISI = 3.56) are significantly increased compared to before correction (Wilcoxon test $p < 0.01$) suggesting the well mixture of cells in each T cell clusters, **C.** Cell count after assigning predicted cell clusters based on the original T cell panel, **D.** Average expression distributions of variable key proteins in each cluster across samples, scaled within each cell cluster.

**A**

scRNA-seq of human T cells in blood (60,725 cells)

↓ gating by CD4+CD8A-CD8B-CCR7-

CCR2+ CD4 (11 cells) vs CCR2- CD4 (336 cells)

↓

338 genes were differentially expressed (adjusted p-value<0.05)

**B** **C**

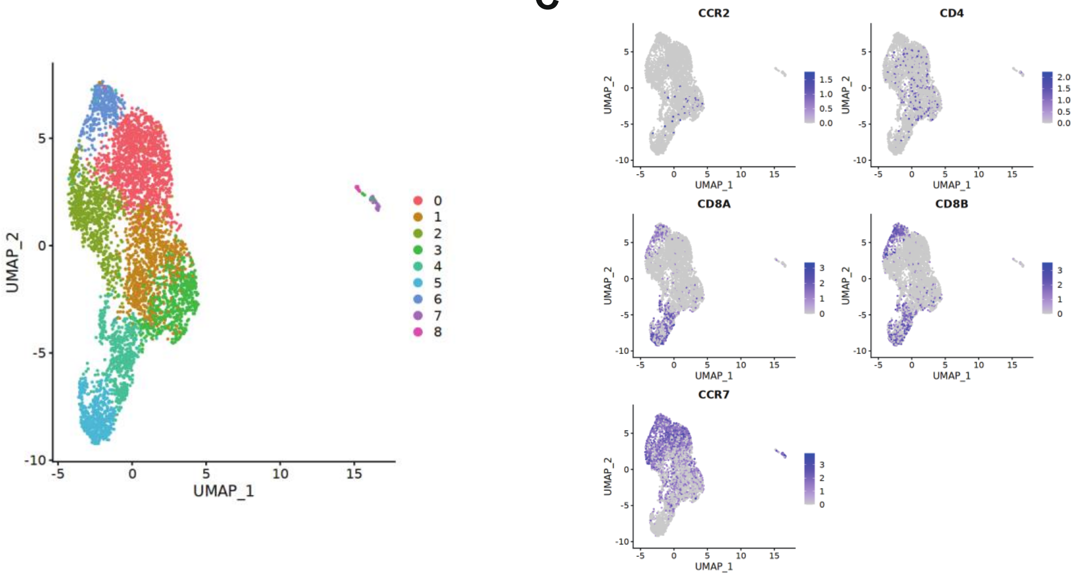


**Extended Data Fig.9: Meta-analysis a single-cell RNA-seq blood dataset enriching CD3+ T cells to infer gene signatures of CCR2+CD4+ T cells. A**. Workflow and results focusing on CCR2+ vs CCR2- within CD4+ T cells using data from GSM3589419 in GSE126030, **B**. UMAP colored by fine-grained cell states in the single-cell RNA -sequencing data, **C**. Expression of selected genes of interest.

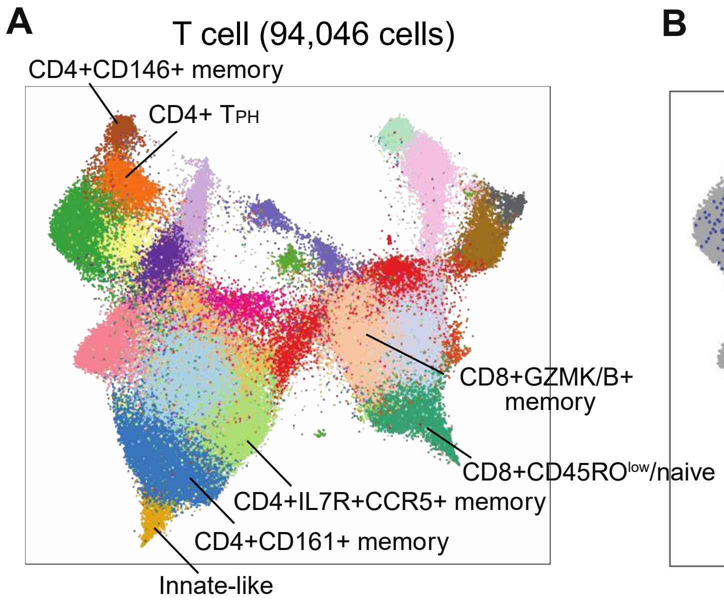


**Extended Data Fig.10: Expression of *CCR2* mRNA in the synovium of RA patients[37]**. **A.** T cell clusters identified in the synovium of RA patients in the UMAP space. The annotations of *CCR2*-expressing clusters are labeled. **B.** *CCR2*-expressing cells in the UMAP. Expressing cells are colored in blue.

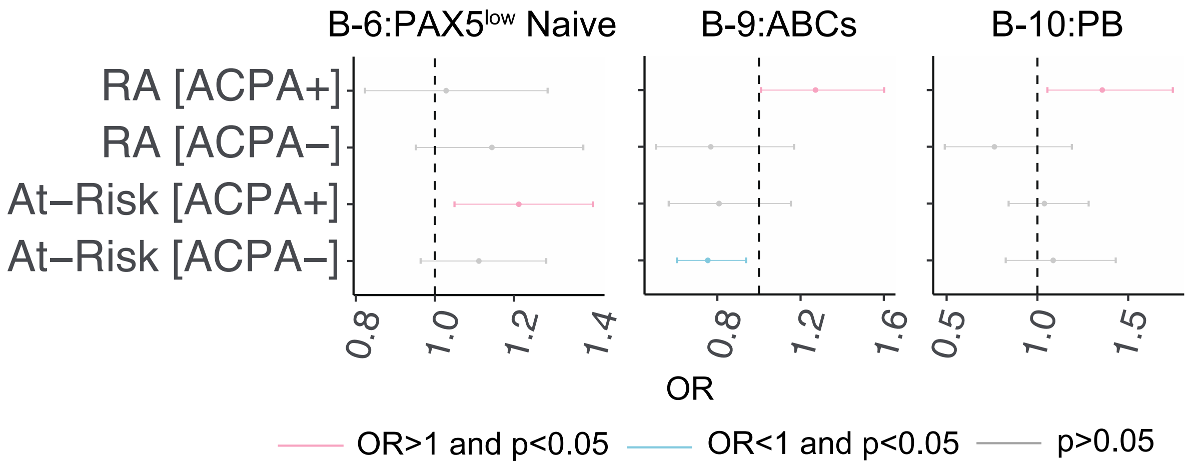


**Extended Data Fig.11: Association with each clinical subgroup according to ACPA status in At-Risk and RA (vs controls) for selected B cell populations.** Error bars represent 95% confidence intervals for the odds ratio. Statistical results are obtained by adjusting for age and sex.

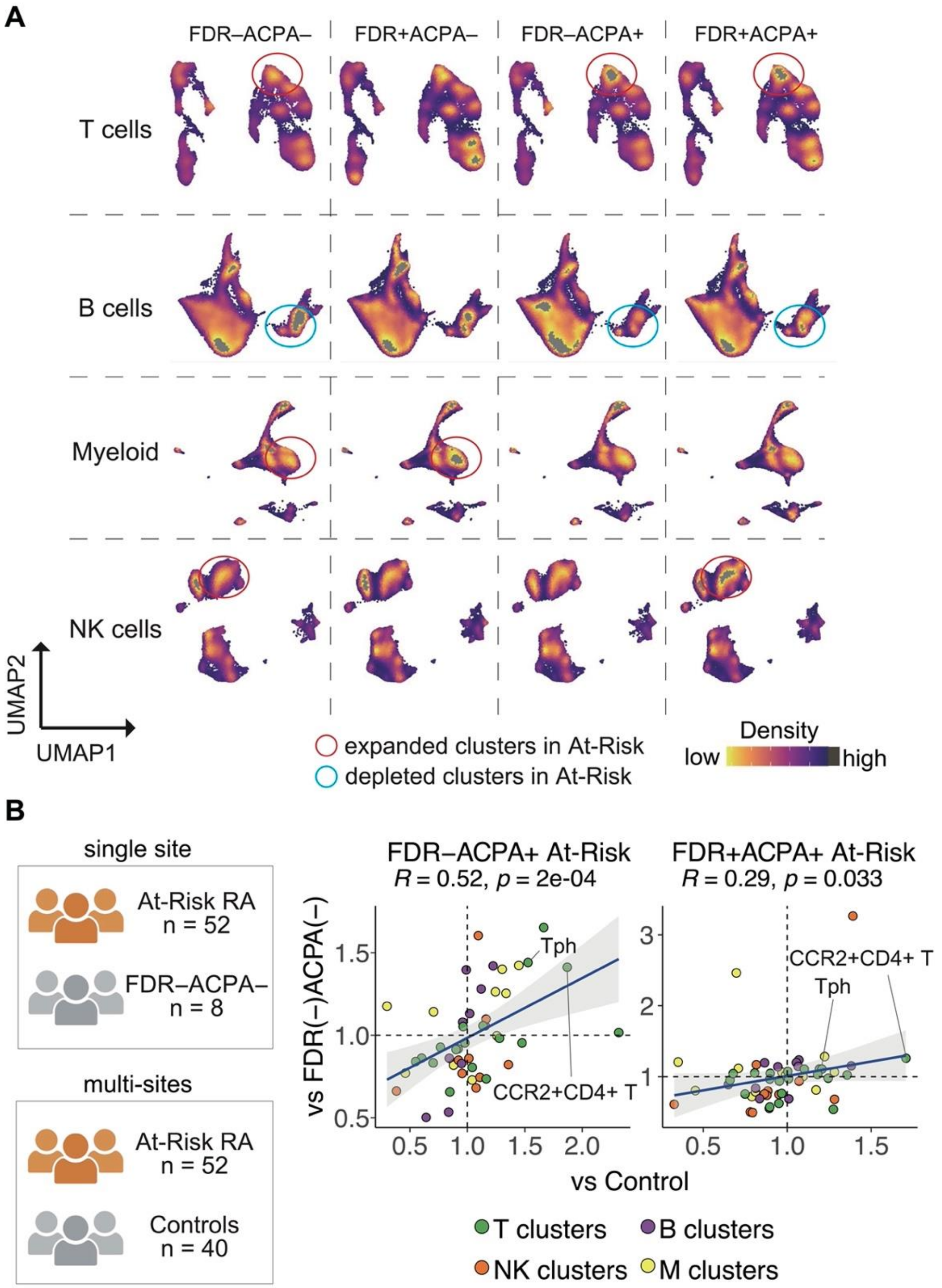


**Extended Data Fig.12: Sensitivity analysis for different control groups from multi-clinical sites. A**. Density plot by family history and ACPA status according to cell types, **B**. Correlation plot of odds ratios comparing At-Risk RA subgroups with FDR−ACPA− controls (y-axis, n=8) from the SERA cohort (y-axis) or healthy controls from other clinical sites (x-axis, n=40). Dots are colored by immune cell types. Of total association tests, 77 cell clusters were included; outliers of the odds ratio (top 99%ile and bottom 1%ile) or size of clusters are lower than 25%ile among all clusters, and results with infinite confidence intervals for the odds ratio were excluded. Statistical results are adjusted for age and sex. Correlation coefficients and p-values were obtained by Spearman's correlation test.
